# RNA Polymerase II hypertranscription at histone genes in cancer FFPE samples

**DOI:** 10.1101/2024.02.28.582647

**Authors:** Steven Henikoff, Ye Zheng, Ronald M. Paranal, Yiling Xu, Jacob E. Greene, Jorja G. Henikoff, Zachary R. Russell, Frank Szulzewsky, H. Nayanga Thirimanne, Sita Kugel, Eric C. Holland, Kami Ahmad

**Affiliations:** Basic Science Division, Fred Hutchinson Cancer Center, Seattle, WA, USA; Howard Hughes Medical Institute, Chevy Chase, MD, USA; Present address: Department of Bioinformatics and Computational Biology, University of Texas MD Anderson Cancer Center, USA; Human Biology Division, Fred Hutchinson Cancer Center, Seattle, WA, USA; Molecular Medicine and Mechanisms of Disease PhD Program, University of Washington, Seattle, WA, USA

**Keywords:** Gene Regulation, Epigenomics, HER2 amplification, Mitochondrial DNA, Meningioma, Whole-arm aneuploidy, Centromeres

## Abstract

Genome-wide hypertranscription is common in human cancer and predicts poor prognosis. To understand how hypertranscription might drive cancer, we applied our FFPE-CUTAC method for mapping RNA Polymerase II (RNAPII) genome-wide in formalin-fixed paraffin-embedded (FFPE) sections. We demonstrate global RNAPII elevations in mouse gliomas and assorted human tumors in small clinical samples and discover regional elevations corresponding to *de novo* HER2 amplifications punctuated by likely selective sweeps. RNAPII occupancy at replication-coupled histone genes correlated with WHO grade in meningiomas, accurately predicted rapid recurrence, and corresponded to whole-arm chromosome losses. Elevated RNAPII at histone genes in meningiomas and diverse breast cancers is consistent with histone production being rate-limiting for S-phase progression and histone gene hypertranscription driving overproliferation and aneuploidy in cancer, with general implications for precision oncology.

## Introduction

Upregulation or amplification of oncogenic transcription factors is common in cancer. For example, mis-regulation of the MYC transcription factor has been observed in most human cancers (*1*), though exactly how increased MYC binding to gene regulatory elements drives cancer has been controversial (*2, 3*). More generally, a global increase in transcriptional output, or hypertranscription, is associated with poor prognosis (*4, 5*). However, it is difficult to reconcile promiscuous incremental increases in expression of thousands of genes with the presumed direct action of oncogenic transcription factors in activating expression of target genes to drive tissue-specific malignancies. Alternatively, hypertranscription in cancer may be relevant only to the subset of genes producing protein products that are rate-limiting for proliferation. For example, the multi-subunit enzyme, ribonucleotide reductase (RNR) – which is required for converting RNA bases to DNA bases – is rate-limiting for DNA synthesis, and RNR activity is a target of widely used anti-cancer catalytic inhibitors (*6*). Similarly, chromatin components such as histones are rate-limiting for proliferation (*7–9*), insofar as all newly synthesized DNA must be packaged into nucleosomes every cell cycle, and the 64 genes encoding all 5 histone subunits must produce ∼5% of a human cell’s total protein during S-phase (*10*). RNAPII is so abundant over histone genes during replication that S-phase-specific global depletion of transcription has been observed in mouse embryonic stem cells (*9*). Replication-coupled (RC) histone mRNAs are not polyadenylated and so are essentially absent from RNA-seq libraries, and the possibility that they drive over-proliferation has been overlooked.

To explore hypertranscription in cancer proliferation, we apply Cleavage Under Targeted Accessible Chromatin (CUTAC) in mouse and human formalin-fixed paraffin-embedded (FFPE) tumor samples (*11*). We document hypertranscription using antibodies to RNA Polymerase II (RNAPII) in three of four mouse brain tumors and three of seven diverse human tumors. In two other human tumors our method discovered amplifications of the HER2 gene region (*12*) and likely selective sweeps (*13*) around other genes nearby. We also found that RNAPII hypertranscription was especially prominent at RC histone genes, leading us to ask whether hypertranscription in cancer is an adaptation to produce more histones at S-phase for more rapid replication. As a critical test of our hypothesis, we performed FFPE-CUTAC on an unselected set of human meningiomas, which are mostly benign human brain tumors that infrequently recur as malignant tumors. We found that FFPE-CUTAC RNAPII occupancy at the 64 RC histone genes predicted WHO grade better than other biomarkers, with similar results for malignant breast tumors. Integration of FFPE-CUTAC data with existing RNA-seq data from 1298 meningiomas accurately separated malignant from benign tumors in predicting recurrence. We also observed striking correlations between levels of RNAPII at histone genes and the number whole-arm chromosome losses in both meningiomas and breast tumors. Successful prediction of tumor aggressiveness based on RNAPII occupancy at histone genes establishes an unanticipated cancer driver paradigm that may also lead to the generation of whole-arm aneuploidies, while opening up diagnostic possibilities not previously considered.

## Results

### RNAPII Hypertranscription varies between tumors

We recently developed FFPE-CUTAC to directly map RNAPII in fixed tissue samples (*11*). This provides a DNA-based method for measuring gene expression, instead of RNA-based methods that are limited by RNA instability and variable transcript half-lives. We assessed RNAPII across candidate *cis-*regulatory elements (cCREs) defined by the ENCODE project, which includes genes and regulatory elements. We compared normal mouse brain tissue to adjacent tumors induced by different transgene drivers: a ZFTA-RELA (RELA) transcription factor gene fusion driving an ependymoma (*14*), a YAP1-FAM118b (YAP1) transcriptional co-activator gene fusion driving an ependymoma (*15*), and overexpression of the PDGFB tyrosine-kinase receptor ligand driving a glioma (*16*). We observed that significantly upregulated cCREs were more frequent than downregulated cCREs (**fig. S1**). To sensitively detect RNAPII hypertranscription (**Fig. 1A**), where the absolute change is important, we first counted the number of mapped fragments spanning each base-pair in a cCRE scaled to the mouse genome coverage and averaged the normalized counts over that cCRE. We then plotted Tumor minus Normal (T-N) counts on the *y*-axis versus the average RNAPII signal (*17*) on a log_10_ scale on the *x*-axis for clarity. This revealed clear hypertranscription (Tumor >> Normal RNAPII) for the RELA tumor (**Fig. 1B**). Two PDGFB-driven tumors differed in hypertranscription, high in PDGFB-1 (**Fig. 1C**) and very low in PDGFB-2 (**Fig. 1D**), whereas the YAP1 tumor showed weak hypotranscription (**Fig. 1E**). To determine whether hypertranscription is specific to any particular class of regulatory element(s), we divided the data into the five ENCODE-annotated cCRE categories: Promoters (24,114), H3K4me3-marked cCREs (10,538), Proximal Enhancers (108,474), Distal Enhancers (211,185) and CTCF cCREs (24,072). We observed that the five RNAPII hypertranscription profiles are highly consistent with one another (**fig. S2**), which suggests that RNAPII abundance differences between tumors and normal brains affect all regulatory element classes.

**Figure 1:**
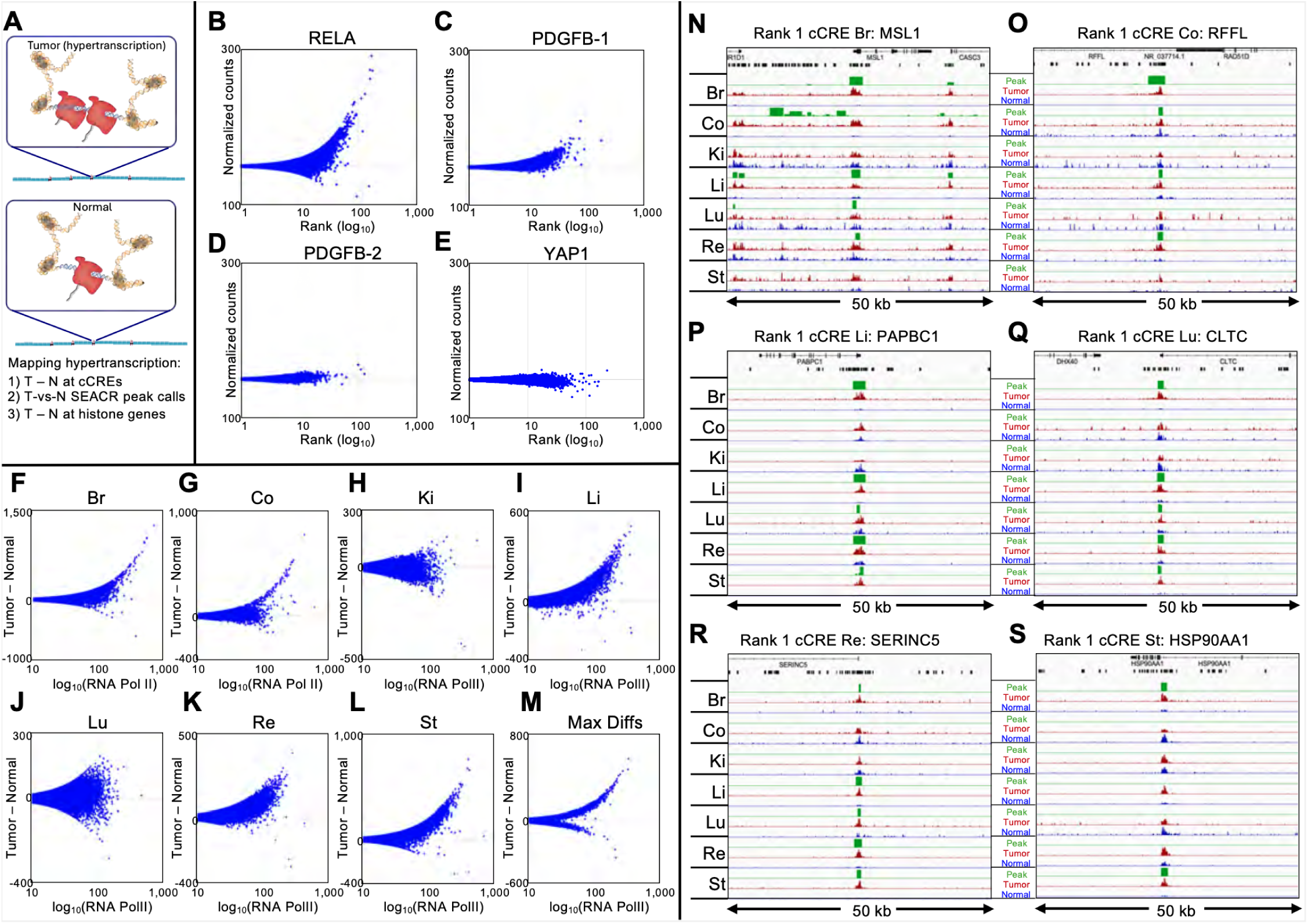
RNAPII-Ser5p FFPE-CUTAC directly maps hypertranscription. (**A**) Model for hypertranscription in cancer: Paused RNA Polymerase II (RNAPII) at active gene regulatory elements such as promoters and enhancers in-creases on average over the cell cycle, resulting in a net proportional gain in RNAPII occupancy across the genome. Using RNAPII FFPE-CUTAC we can map hypertranscription genome-wide using three complementary approaches: 1) Ge-nome-scaled Tumor (T) minus Normal (N) counts at cCREs, 2) T – N at replication-coupled histone genes and 3) SEACR Tumor peak calls using Normal as the background control. (**B-E**) T – N versus log(T + N)/2 plots showing hypertranscrip-tion mapped over the 343,731 annotated mouse cCREs for tumor and normal sections dissected post-tagmentation from a 10 micron FFPE slice from each of four different paraffin blocks. Hypertranscription of a cCRE is defined as the excess of RNAPII-Ser5p in the indicated tumor over normal (Tumor minus Normal in normalized count units for Mm10-mapped fragments pooled from the same slide). (**F-M**) All fragments were pooled from four slides from the same paraffin block and the number of fragments equalized between tumor and normal for each of the seven cancers. T – N versus log(T + N)/2 plots showing hypertranscription mapped over the 984,834 annotated mouse cCREs for tumor and matched normal sections from 5 micron FFPE slices. Max Diffs displays the Tumor minus Normal maximum of the seven samples for each cCRE. (**N-O**) For each of the indicated tumors, tracks are shown for 50-kb regions around the #1-ranked cCRE based on Tumor (dark red) and Normal (blue) counts. Raw data tracks were group-autoscaled together for tumor (red) and normal (blue), where SEACR Tumor peak calls (green) use Normal as the negative control. Gene annotations and cCREs (black rectangles) are shown at the top.

To expand on our findings of RNAPII hypertranscription based on transgene-driven mouse brain tumors to a diverse sample of naturally occurring cancers, we obtained 5-µm FFPE sections on slides prepared from paraffin blocks of anonymous human tumor and matched normal sections from the same patient (**fig. S3**). We performed FFPE-CUTAC and rank-ordered each pair by Tumor minus Normal (T-N) differences to test for RNAPII hypertranscription based on the 984,834 ENCODE-annotated human cCREs. We observed clear hypertranscription in five of the seven tumors (breast, colon, liver, rectum and stomach) and for the composite of all samples (**Fig. 1F-M, fig. S4**). In contrast, the kidney and lung tumor samples tested showed essentially no hypertranscription, which implies that hypertranscription is a common, but not a defining feature of cancer (*2, 4*).

We also applied a computational method independent of annotations to assess RNAPII hypertranscription in FFPE-CUTAC profiles of patient samples. The SEACR (Sparse Enrichment Analysis for CUT&RUN) tool is specifically designed for low read-count data (*18*). To customize SEACR for hypertranscription in cancer, we replaced the background control with the normal sample in each pair, merged fragment data, removed duplicates and equalized read numbers for our seven human Tumor/Normal pairs. SEACR reported a median of 4,483 peaks that were elevated in tumors, whereas when Tumor and Normal were exchanged only a median of 15 peaks elevated in normal tissue were identified, demonstrating that RNAPII hypertranscription is more common than RNAPII hypotranscription in these cancer samples.

We next asked whether SEACR Tumor-versus-Normal peak calls corresponded to the 100 top cCREs ranked by T-N in the overall list representing all seven human tumors. Remarkably, all 100 cCREs at least partially overlapped one or more SEACR Tumor/Normal peak calls, and in addition, the large majority of the 100 top-ranked cCREs intersected with overlapping SEACR peak calls from multiple Tumor/Normal pairs (**Table S1**). Each of the #1-ranked cCREs in the breast, colon, liver, lung and rectum tumor samples respectively intersected MSL1, RFFL, PABPC1, CLTC and SERINC5 genes and overlapped SEACR peak calls in 4-5 of the 7 tumors (**Fig. 1N-O, fig. S5**). On average, the same cCRE overlapped SEACR/Normal peak calls in 3.7 of the 7 tumors (**Table S1**). No SEACR peaks were observed for the kidney sample, as expected given the lack of detectable RNAPII hypertranscription over cCREs. We conclude that the most strongly RNAPII-hy-pertranscribed regulatory elements tend to be strongly hypertranscribed in multiple human cancers of different types, including liver cancers from different individuals (**fig. S6**).

Interestingly, we observed a much lower level of mito-chondrial DNA (mtDNA) in most tumor samples than in their matched normal samples for both mouse and human (**fig. S7A-B**), suggesting that these tumors contain fewer mitochondria. To test this interpretation, we mined publicly available ATAC-seq data from both the TCGA and ENCODE projects and observed sim-ilar reductions in cancer samples (**fig. S7C-D**). Such reductions in mtDNA have been reported based on whole-genome sequencing (*19*).

### HER2 amplifications with linkage disequilibrium in human tumors

All but one of the top 25 cCREs are located on Chro-mosome 17 (**Table S1**). Eight of these cCREs are with-in Chr17q12 and 13 are within Chr17q21, each span-ning a few hundred kilobases in length in the breast tumor sample not seen in the normal tissue (**Fig. 2A-B, fig. S8**). For the colon tumor sample, a broad re-gion of RNAPII enrichment is sharply defined within Chr17q21. High RNAPII occupancy over the cCREs in Chr17q12-21 can account for most of the RNAPII hypertranscription signal in the breast and colon sam-ples, centered over the ERBB2 gene (**fig. S9**). ERBB2 encodes Human Epidermal Growth Factor Receptor 2 (HER2), and is commonly amplified in breast and other tumors and is a target of therapy (*12*). As our measures of RNAPII hypertranscription are normalized with re-spect to the human genome coverage, amplification of a region will appear as a proportional increase in the level of FFPE-CUTAC signal, so that we can interpret regional RNAPII hypertranscription in the breast and colon tumor samples as revealing amplification events distinct from the global upregulation that defines hyper-transcription.

**Figure 2:**
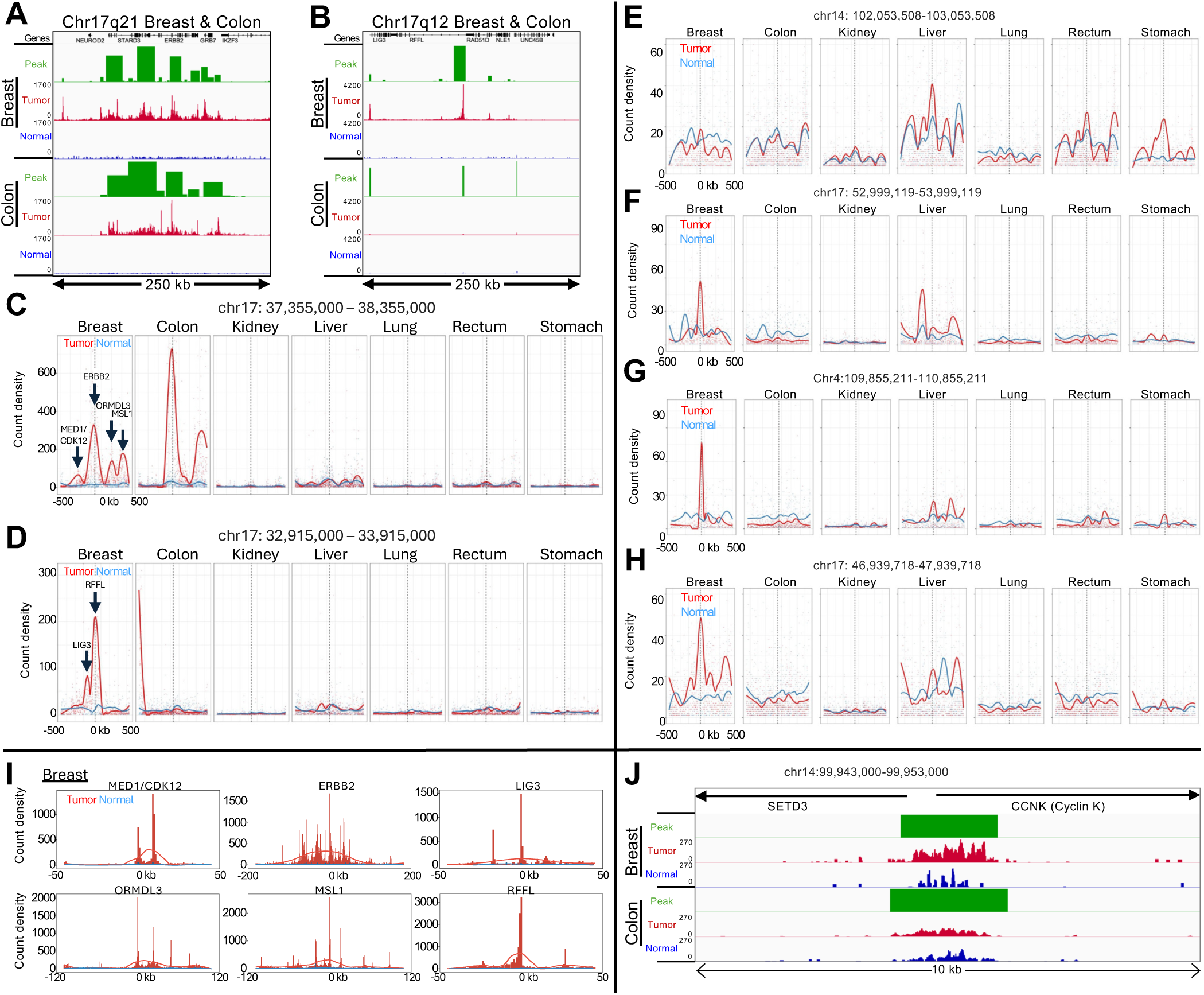
RNAPII levels identify likely HER2 amplifications and regions of linkage disequilibrium. (**A-B**) Normalized count tracks and SEACR peak calls for the 250-kb region on Chromosome 17q21 (A) and for the 250-kb 17q12 region (B) amplified in the breast tumor but evidently not in the colon tumor. Tracks were group-autoscaled together for Tumor (red) and Normal (blue), where SEACR Tumor peak calls (green) use Normal as the negative control. Broad regions of prominent RNAPII hypertranscription indicate likely HER2 amplifications in both tumors. (**C-D**) Bins of 1-kb were tiled over each 1 Mb region centered on the highest peak in Chr17q21 (C) corresponding to the ERBB2 promoter in Chr17q21, and the RFFL promoter in Chr17q12 (D), and count density within each bin is plotted with curve-fitting and smoothing. Each of the six summits in the Breast tumor sample is centered over the promoter peak indicated by an arrow. (**E-H**) Same as (C-D) for top-ranked loci outside of the HER2 amplicon. (**I**) Individual broad summits in (D-E) were zoomed-in and rescaled on the *x*-axis centered over the indicated promoter peak and superimposed over data tracks scaled to the height of the central peak. (**J**) Data tracks for the *CCNK* promoter region, where the normalized count increase in the Breast tumor relative to normal over the 10-kb region shown is 5.4-fold and for Colon is 2.1-fold. The range for the other five tumors is 0.9-2.5.

To confirm that the broad regions of RNAPII enrichment around the ERBB2 promoter correspond to HER2 amplifications in the Breast and Colon patient samples, we applied SEACR broad-peak calling, which densely tiled a ∼150 kb region centered over the ERBB2 promoter (**Fig. 2A-B**). To ascertain whether dense tiling using SEACR can detect amplification events, we called SEACR broad peaks on our published K562 RNAPII-Ser5p CUTAC datasets (*20, 21*). When we examined Chromosome 22q, we observed a single region heavily tiled with broad SEACR peaks corresponding to an annotated amplification specific for K562 cells. Zooming in revealed that the end of the densely tiled region corresponded precisely to the t(9;22)(q34;q11) translocation breakpoint of BCR-ABL in BCR (**fig. S10A-B)**, which we confirmed by observing a broad SEACR peak on the ABL1 side of the translocation breakpoint (**fig. S10C**). Thus, our approach using RNAPII-Ser5p CUTAC and SEACR can identify and precisely map regional amplifications such as are found in tumors with BCR-ABL and HER2 amplifications.

To delineate possible RNAPII hypertranscription features within Chr17q12 and Chr17q21, we binned successive 1-kb tiles over each 1 Mb region centered on the highest peak, corresponding to the ERBB2 promoter in Chr17q21 and the RFFL promoter in Chr17q12, and plotted count density within each bin with curve-fitting and smoothing. Remarkably, multiple broad summits appeared in both breast and colon tumor-ver-sus-normal tracks, and the six summits in the breast tumor sample accounted for the six highest ranked Chr17 promoter peaks (**Fig. 2C-D**). We similarly plotted count densities of the four highest ranking cCREs outside of Chr17q12-21 (**Table S1**), but tumor peaks in these regions were at least an order-of-magnitude lower than the ERBB2 peaks in the breast and colon tumor samples (**Fig. 2E-H**). Of the six summits in the breast tumor sample, ERBB2 and MSL1 also ap-peared in the colon tumor sample, whereas no other samples showed prominent summits above normal in Chr17q12-21 (**Fig. 2C-D**). MSL1 encodes a subunit of a histone H4-lysine-16 acetyltransferase complex required for upregulation of the mammalian X chromosome (*22*).

We next superimposed each of the six summits in the Chr17q12-21 region in the breast tumor sample over the genomic tracks on expanded scales for clarity, centered over the highest promoter peak in the region (**Fig. 2I**). For ERBB2, the ∼100 kb broad summit is almost precisely centered over the ∼1 kb wide ERBB2 promoter peak. Although the other summits are less broad, each is similarly centered over a promoter peak. Insofar as there are multiple summits much broader than the promoter peaks that they are centered over, our results are inconsistent with independent upregulation of promoters over the HER2-amplified regions. Rather, it appears that a HER2 amplification event was followed by clonal selection for broad regions around ERBB2 and other loci within each amplicon, consistent with the observation of clonally heterogeneous HER2 amplifications in primary breast tumors by whole-genome sequencing (*23*). Clonal selection may be driven by selective sweeps (*24*) following amplification events that generate extrachromosomal DNA in double-minute acentric chromosomes, which partition unequally during each cell division (*13, 25, 26*). Such copy number gains within a tumor can result in intra-tumor heterogeneity (*26, 27*) and are potential factors for resistance to therapy (*28*). FFPE-CUTAC thus potentially provides a general diagnostic strategy for detection and analysis of amplifications and clonal selection during cancer progression and therapeutic treatment.

One of the summits in the breast tumor sample absent from the colon tumor sample corresponds to the bidirectional promoters of MED1 and CDK12, both of which have been shown to functionally cooperate with co-amplified ERBB2 in aggressive breast cancer (*29, 30*). MED1 encodes a subunit of the 26-subunit Mediator complex, which regulates RNAPII pause release, and CDK12 is the catalytic subunit of the CDK12/Cyclin K kinase heterodimer complex, which phosphorylates RNAPII for productive transcriptional elongation (*31, 32*). We wondered whether the co-amplification of these RNAPII regulators might contribute to hyper-transcription in this tumor. As Cyclin K is the regulatory subunit of the CDK12 kinase, we would expect that the *CCNK* gene that encodes Cyclin K would be strongly upregulated in the breast tumor but not necessarily in the colon tumor. Indeed, we saw a 5.4-fold increase in RNAPII-S5p over the *CCNK* promoter in the breast tu-mor relative to adjacent normal tissue, whereas in the colon tumor there is a 2.1-fold increase (**Fig. 2J**), con-sistent with RNAPII hypertranscription directly driven in part by CDK12 amplification in this particular patient’s tumor.

### RNAPII over histone genes predicts aggressive-ness in meningiomas and breast tumors

To evaluate how effectively FFPE-CUTAC can resolve differences between the seven tumor samples, we con-structed a cCRE-based UMAP including all 114 individ-ual human datasets with >100,000 mapped fragments (median 925,820). Whereas normal samples produced mixed clusters, tumor samples formed tight homoge-neous clusters separated by tissue type (**Fig. 3A**). This implies that paused RNAPII at regulatory elements is more discriminating between tumors than between the tissues that the tumors emerge from. Relatively few samples and shallow sequencing depths were need-ed for tight clustering; for example, the stomach tumor cluster comprised four samples from four different ex-periments with a median of ∼470,000 mapped frag-ments (**Fig. 3B**). The colon and breast tumor samples, which share HER2 amplifications, clustered immedi-ately adjacent to one another.

**Figure 3:**
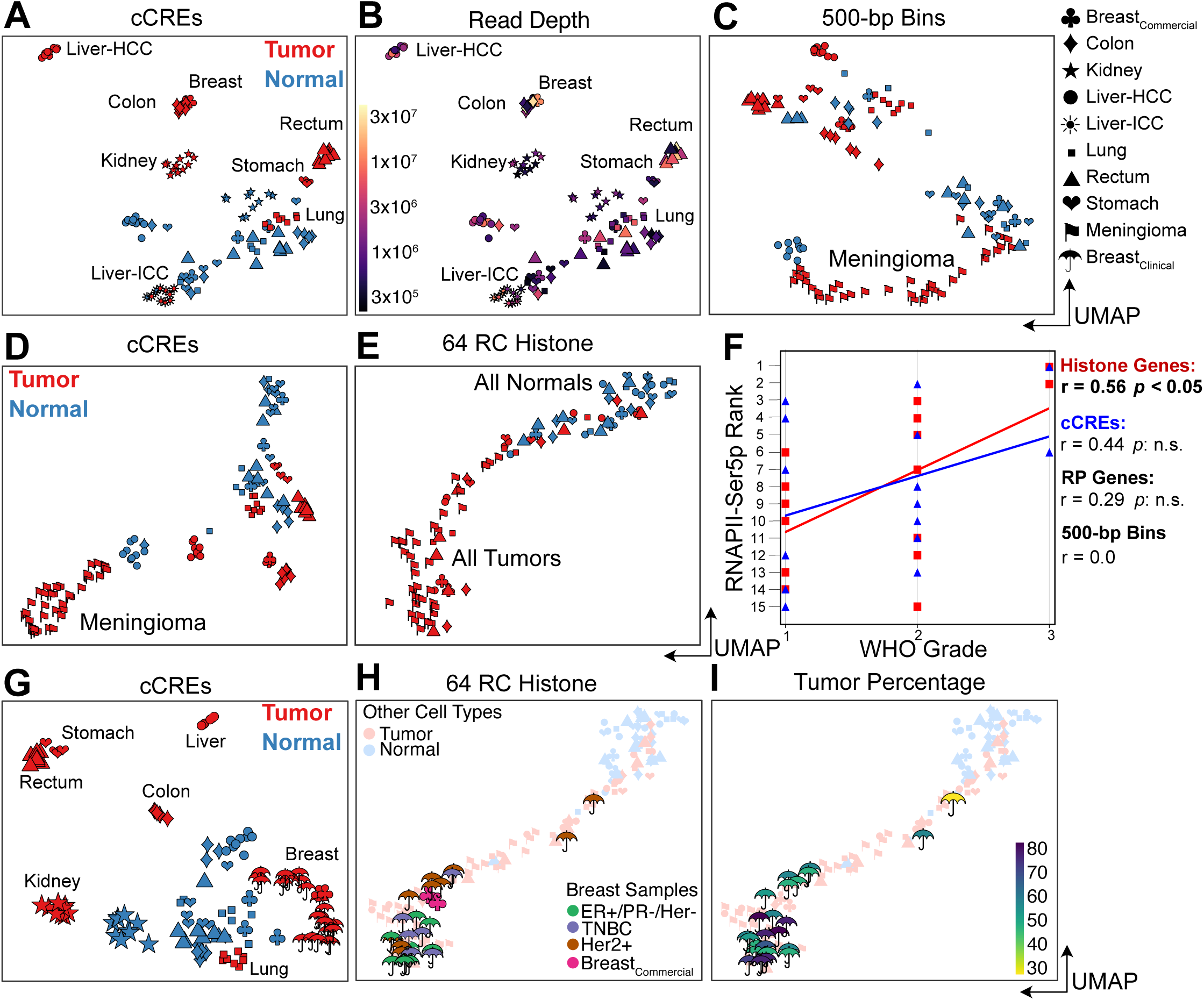
RNAPII over histone genes correlates with aggressiveness in meningiomas (A-F) and breast tumors (G-I) **(A)** UMAP of 114 human tumor samples and replicates including those from 30 meningiomas. HCC: Hepatocellular carci-noma; ICC: Intrahepatic cholangiocarcinoma. (**B**) Same as (A) colored for sequencing depth and indicating homogeneous tumor clusters. (**C**) UMAP of RNAPII FFPE-CUTAC data based on 500-bp bins over the human genome. (**D**) Same as (C) for cCREs. (**E**) Same as (C) for the 64 Replication-Coupled (RC) histone genes. (**F**) WHO grade correlates best with RNAPII occupancy over histone genes. (**G**) Same as (A) excluding meningiomas but including samples and replicates from 13 additional breast tumors. (**H**) Same as (E) for breast tumor samples and replicates, colored for tumor type. (**I**) Same as (H) colored for tumor percentage.

To verify that these differences in global upregulation of cCREs are related to tumor growth, we examined the profiles of the RC histone genes. As these exceptionally S-phase-dependent histone loci are expressed in proportion to the amount of replicated DNA (*7*), we wondered whether cancer cell hypertranscription functions to increase engaged RNAPII at these loci to load up on histones at S-phase for more rapid cell proliferation. For both the 64 mouse and the 64 human annotated replication-coupled genes we observed differences between tumor samples consistent with RNAPII hypertranscription at cCREs differing between samples (**fig. S11**). We reasoned that if histone production at S-phase is rate-limiting for proliferation, then we would observe a correlation between cancer aggressiveness and RNAPII, specifically over histone genes. As a test of this hypothesis, we applied FFPE-CUTAC to 4-micron FFPEs from 30 meningioma patients and constructed a UMAP that also included the Tumor-Normal pairs described above. Using RNAPII abundance within 500-bp bins for UMAP construction, we found that the meningiomas clustered separately from other tumors and from normal, and using RNAPII abundance on the cCREs, they formed a single tight cluster (**Fig. 3C**). When we performed the same UMAP construction using only RNAPII abundance over the RC histone genes, we found that all tumors regardless of type clustered together in a cline that overlapped normal at one end (**Fig. 3D**). This suggested that the RNAPII at histone genes can distinguish cancer from normal but is insensitive to cancer type differences. In a double-blind test of whether the cline overlapping normal in 2D UMAP space is an indication of tumor aggressiveness, we rank-ordered the 15 samples with WHO grades based on distance from normal samples for RNAPII occupancy in 500-bp bins, cCREs, histone genes or ribosomal protein genes (**Fig. 3E**). Best performance was for the RC histone genes (r=0.57, *p*<0.05, **Fig. 3F**), consistent with our hypothesis that high RNAPII levels at histone genes drive proliferation in cancer.

To determine whether RC histone genes can predict cancer aggressiveness in an invasive tumor type, we performed FFPE-CUTAC on a set of 10-µm breast tumor FFPEs from 13 patients representing three major subtypes. When we included these samples with the 7 diverse tumor and normal samples (**Fig. 1F-L**) and used cCREs for UMAP construction, we observed tight clustering strictly according to tumor type (**Fig. 3G**). However, when we used RC histone genes for UMAP construction, we observed a single large cluster at one end of a cline that overlapped normal at the other end (**Fig. 3H-I**). In total, 24 of the 26 individually amplified samples were entirely within the single large tumor-only cluster. We conclude that RC histone genes alone can predict cancer aggressiveness in both non-invasive meningiomas and multiple invasive breast cancer subtypes.

### RNAPII over histone genes accurately predicts rapid recurrence in meningiomas

WHO grade is a coarse predictor of recurrence (*33*), so to predict rapid recurrence for each tumor sample, we integrated FFPE-CUTAC data with RNA-seq data to use nearest neighbors on a UMAP constructed from available RNA-seq data. We defined a gene as spanning from the 3’-most transcript end through the 5’-most end, stopping when either end of the next gene or LINE element is reached. Based on normalized counts over each RefSeq-annotated human gene, we successfully integrated the public frozen meningioma RNA-seq samples with our 30 meningioma FFPE-CUTAC samples (**fig. S12**). Remarkably, 17 of 19 matching samples from the same meningioma patient showed near-coincidence of frozen RNA-seq and FFPE-CUTAC.

To predict patient clinical outcomes, we employed a data-driven strategy to classify FFPE-CUTAC samples with overall RNAPII signal at histone genes and leveraged the top 20 shared nearest neighbors of RNA-seq samples to obtain the meningioma patient recurrence information (**fig. S13A**). Indeed, we found a highly significant association between high histone signals in FFPE-CUTAC samples and rapid patient recurrence (**fig. S13B**). Using only the normalized FFPE-CUTAC counts at the 64 histone genes, we could distinctly separate the five most rapidly recurring from the 25 other tumors with high significance (*p*<10^-8^, **Fig. 4A-B** left). This observation aligns with the generally low recurrence rate of meningiomas, which are predominantly benign (*34*). In contrast, levels of RNAPII over Ribosomal Protein genes failed to significantly separate rapidly recurring from benign, regardless of the thresholds applied to designate high Ribosomal Protein gene signal samples (**Fig. 4A-B, fig. S13C**). We also applied the same logic to determine whether reduced mtDNA is a feature of malignant meningioma but observed no significant difference (**Fig. 4A-B, fig. S13D**). Although we did observe significant separation of rapidly recurring from benign using Chr22q, the most frequently lost whole-arm (*33*) (**Fig. 4A-B** right), and significant sepa-ration for Chr1q gains and Chr6p losses (**fig. S13E-F)**, the levels of significance were much lower than for separation using RNAPII at the RC histone genes. Our finding that high levels of RNAPII over RC histone genes accurately predicts poor outcomes in meningio-ma could imply a causal basis.

**Figure 4:**
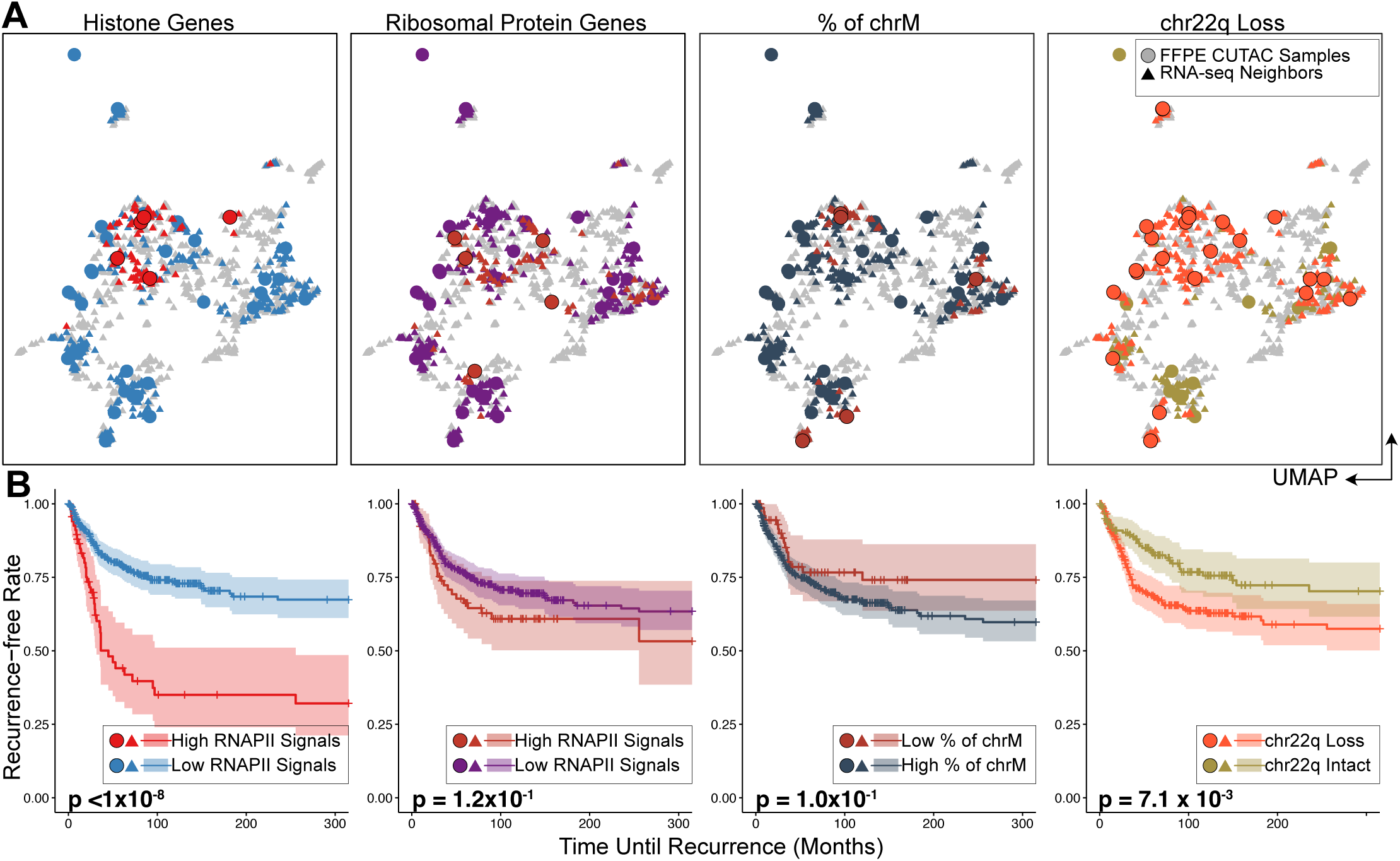
RNAPII over histone genes uniquely predicts recurrence in meningiomas. (**A**) UMAPs of FFPE-CUTAC and frozen RNA-seq meningioma patient samples grouped by different biomarkers. From the left to right: RNAPII over RC histone genes; RNAPII over ribosomal protein genes; Percent of Chromosome M (chrM = Mitochondrial DNA); Chr22q loss. UMAPs are based on integrating FFPE-CUTAC (circles) with frozen-RNA-seq (triangles) samples (**fig. S10**) using the canonical correlation analysis. For RC histone and ribosomal protein genes, points are colored by high (red) or low (blue for histone genes and purple for ribosomal protein genes) RNAPII enrichment over the corresponding genes for FFPE-CUTAC samples or their shared nearest neighbor RNA-seq samples. For chrM, points are colored by low (red) or high (dark grey) fractions of Chromosome M fragments, with the hypothesis that a low chrM fraction predicts malignant. For chr22q loss, the FFPE-CUTAC samples with chr22q loss and their RNA-seq samples are colored red, while the ones with chr22q intact are colored green. (**B**) Kaplan-Meier (KM) plots of recurrence-free rate as a function of time in months based on each grouping strategy using the biomarkers shown in (C). For fair comparisons between Histone genes, Ribosomal Protein genes and chrM, the same number of samples, i.e., the top five patients, are selected in the malignant group. The fraction of the chrM predictor has settings (6 to 18 samples in the malignant group) with small p values. However, the KM curve order is in the wrong direction for the hypothesis that a low fraction of chrM predicts fast recurrence.

### RNAPII over histone genes predicts whole-arm chromosome losses

As a positive control for separation of malignant from benign, we counted total aneuploidies from RNA-seq data and observed best separation for five malignant and 25 benign tumors, closely matching our prediction based on RNAPII at histone genes over the range of 3-7 predicted malignant (Figure 5A). This correspondence led us to ask whether there is a relationship between overproduction of histones and total aneuploidy by plotting RNAPII levels over the histone genes for each patient as a function of the total number of whole-arm chromosome gains or losses. We observed a weak non-significant positive Spearman correlation with gains (p<0.2), but a highly significant correlation with losses (p<0.006) (**Fig. 5B**). To test whether this excess of losses over gains applied to all chromosome arms, we asked whether for each patient, there was a net increase or decrease in the level of RNAPII over histone genes, where whole-arm gains would show a negative correlation with the number of cCREs lack-ing RNAPII occupancy (no signal over the cCRE) and whole-arm losses would show a positive correlation. Indeed, 38/39 autosomal arms showed a net posi-tive correlation for the meningioma patient population (**Fig. 5C, fig. S14A**), suggesting that RNAPII at his-tone genes predicts whole-arm losses in meningio-ma. We confirmed this excess of whole-arm losses for the breast cancer samples, where we also observed excesses of losses over gains for 38/39 autosomal whole-arm aneuploids (**Fig. 5C, fig. S14B**), as expect-ed if overexpression of histones drives both over-prolif-eration and whole-arm chromosome losses in cancer.

**Figure 5:**
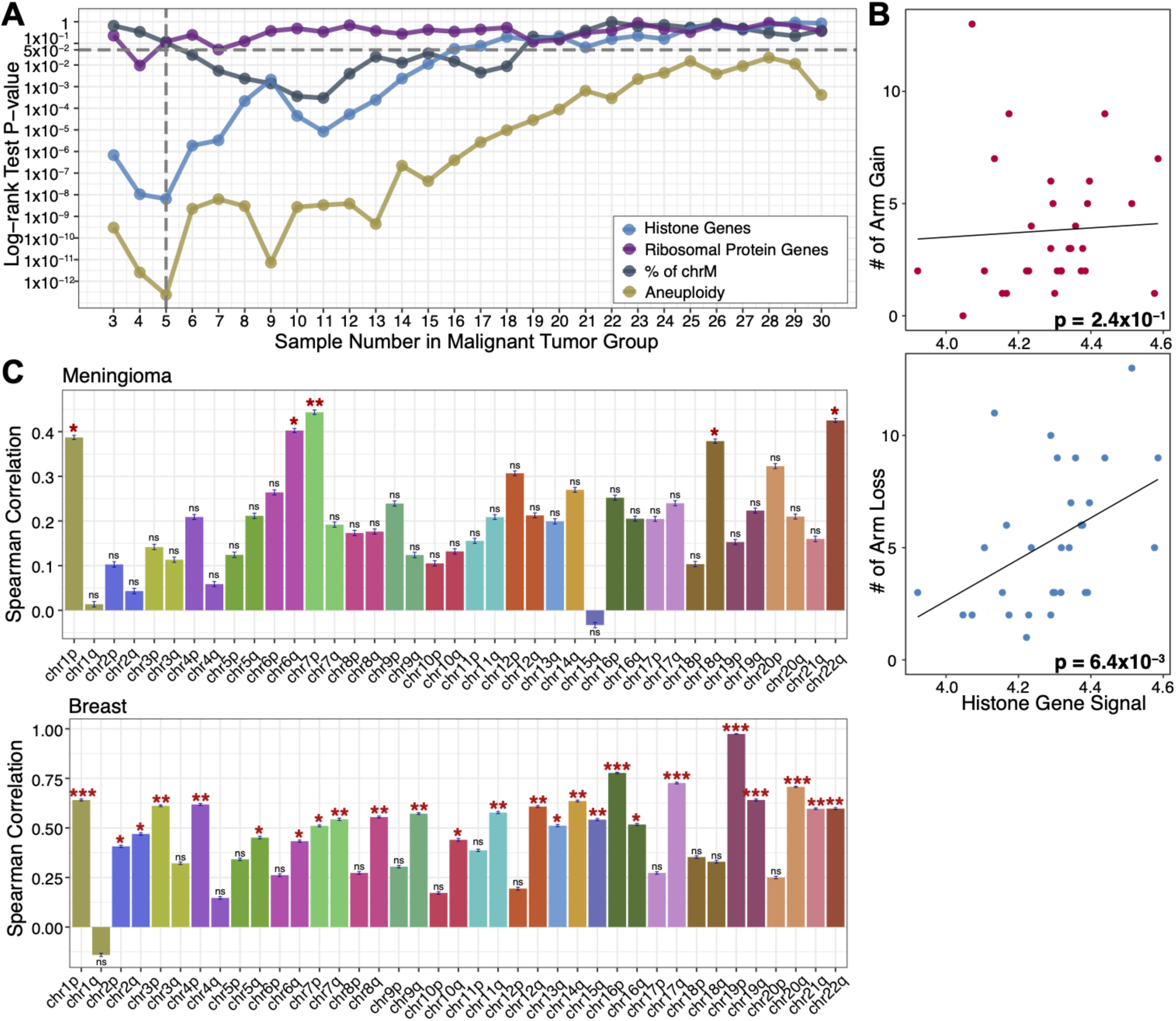
RNAPII over histone genes predicts whole-arm chromosome losses. **(A)** P-values from the log-rank test for the KM separation of malignant from benign for all thresholds of the 30 meningioma patients and the corresponding 36 FFPE-CUTAC samples for Histone genes (blue), Ribosomal protein genes (purple) and % mitochondrial DNA (grey) (**Fig. 4**), also including the combined aneuploidy data for all 39 chromosome arms (gold). (**B**) Scatterplots and Spearman correlations between RNAPII over histone genes and number of whole-arm gains (upper panel) and losses (lower panel) for each of the 30 meningioma patient samples. (**C**) Spearman correlations between the signal for RNAPII over histone genes and null RNAPII occupancies at cCREs for all 39 autosomal arms for Meningiomas (upper panel) and Breast tumors (lower panel). This summarizes correlation coefficients and significances on the correlation scatterplot in fig. S14. A negative correlation is expected if gains are more frequent than losses (i.e., chromosome arm gain will lead to a lower percentage of cCREs regions with zero RNAPII signals) with increasing RC histone gene signal and a positive correlation if losses are more frequent than gains. The error bar describing the standard deviation of the Spearman correlation coefficient is based on 1000 bootstrap estimation. The significance level is indicated by the stars on the top of each bar with *: p-value < 0.05, **: p-value <0.01, and ***: p-value < 0.001.

### Mitotic chromosome segregation errors do not ac-count for loss-over-gain biases

The bias of whole-arm losses over gains in breast tu-mors is far more significant than in meningiomas (**Fig. 5C**), presumably because all of the breast tumors but only ∼1/4 of the meningiomas are malignant. Bias in favor of loss over gain is counter-intuitive in that a 50% loss should be less fit than a 50% gain, as autosomal trisomies occur in ∼0.3% of newborns, but no autoso-mal monosomies survive to term. Whole-chromosome trisomies result from mitotic segregation errors, such as merotelic attachments, extra centrosomes, unattached kinetochores, loss of the spindle assembly checkpoint or cohesion defects. However, whole-arm aneuploid chromosomes are generated by an initial centromere break, and we wondered whether mitotic segregation errors involving broken centromeres could account for the striking biases in favor of losses over gains that we observed. To test this possibility, we took advantage of the two classes of human chromosomes based on the position of the centromere. Whole-arm gains and loss-es produced by centromere breaks are far more fre-quent than focal aneuploidies in cancer (*35*), and using TCGA data we find that whole-chromosome aneu-ploids comprise only ∼15% of the total (**Fig. 6A**). Meta-centric chromosomes have two euchromatic arms and so require a centromere break to generate a whole-arm SCNA (**Fig. 6B**). There are five human acrocentric chromosomes (13, 14, 15, 21 and 22), which have similar kinetochore conformations as metacentrics (*36*), but only a single euchromatic arm. Acrocentrics that are gained or lost by mitotic error will be effectively indistinguishable from those that have undergone a centromere break event. Acrocentric short arms comprise only redundant ribosomal DNA genes and other tandem repeats, and single short-arm gains or losses are not detected in genomic studies. Whole-chromosome meiotic or mitotic segregation errors occur, but centromere position does not predict frequency, based on pre-meiotic mitoses during human oocyte maturation (78% metacentrics expected, 83% observed, n = 52) (*37*). Thus, if mitotic errors are essential for the generation or perpetuation of a significant number of whole-arm SCNAs, then there should be an excess of acrocentrics (both those with and those without cen-tromere breakpoints) relative to metacentrics (all of which must have centromere breakpoints). To distin-guish these models, we have analyzed whole-genome sequencing data from TCGA for acrocentrics and me-tacentrics from 10,674 patients based on allele-specific copy number segmentation analysis across 33 cancer types. However, we observed no significant difference between acrocentrics and metacentrics in the frequen-cy of whole-arm SCNA gains or losses or both (**Fig. 6C, fig. S15**). We conclude that centromere breaks are sufficient to generate the excess of whole-arm losses that we observed.

**Figure 6:**
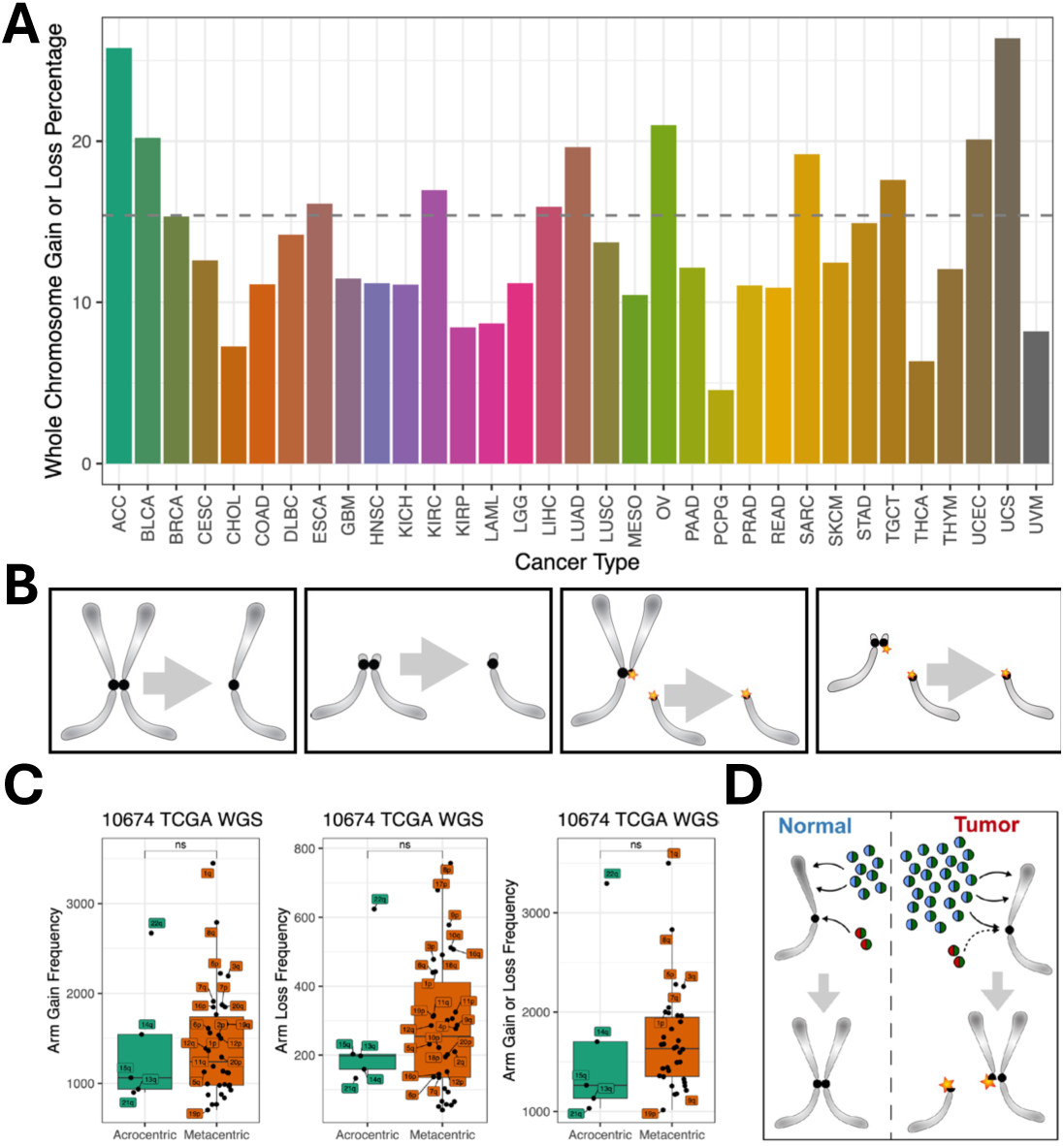
Acrocentric and metacentric whole-arm SCNAs are recovered at similar frequencies in cancer. **(A)** Left to right: A metacentric segregation; An acrocen-tric segregation; A metacentric segregation following a centromere break; An acrocentric segregation following a centromere break. (**B**) Summary of whole-genome sequenc-ing data from the Cancer Genome Atlas (TCGA, https://portal.gdc.cancer.gov/) representing 10,674 cancer patients for 33 cancer types (**fig. S15**). For 32 of 33 cancer types, no significant differences are seen between acrocentrics and metacentrics in the frequencies of whole-arm SCNA gains, losses or both gains and losses (Wilcoxon rank test). Each dot represents a different autosomal chromo-some arm (5 acrocentric long arms and 17 metacentrics). (**C**) We inferred the chromosome arm gain or loss using allele-specific copy number analysis of tumors (<u>ASCAT)</u> (*56*) profiles for each patient. We summed the major and minor alleles of each segment and took the minimum and maximum of the copy number across segments on each chromosome arm. Any increases in the minimal allelic copy number from the diploid copy number of 2 indicate an arm gain. Similarly, any decreases in the maximum allelic copy number from the diploid copy number of 2 indicate a whole-arm loss for the corresponding autosomal arm. (**D**) Model for induction of centromere breaks and aneuploidy by histone gene hypertranscription. Histone H3|H4 dimers (blue-green spheres) are deposited along chromosome arms during DNA replication for chromatin duplication, while CENPA|H4 dimers (red-green spheres) are deposited at centromeres and maintain centromere identity. H3|H4 and CENPA|H4 histone dimers are preferentially deposited (black arrows) at the CAF1 and HJURP assembly factors, respectively. However, the excess H3|H4 dimers produced by hypertranscription in cancer cells compete with CEN-PA|H4 for centromere assembly, reducing centromere function and creating DNA breaks (*45, 46*). Segregation of broken chromosomes leads to whole-arm aneuploidies.

## Discussion

We have demonstrated that elevated RNAPII over genes and regulatory elements is a direct measure of hypertranscription in diverse human cancers and identifies and precisely maps amplifications and selective sweeps in small clinical samples. Likewise, we observed a close correspondence between high RNAPII at the 64 replication-coupled histone genes consistent with cytological evidence of exceptionally high levels of RNAPII at mouse RC histone genes (*9*) and both RNAPII (*7, 8*) and Myc (*38*) at Drosophila histone locus bodies exclusively during S-phase. This led us to hypothesize that the single functional role of hypertranscription in cancer is to produce enough histones to keep up with the requirement for packaging new DNA in cancer cells to proliferate faster than normal cells. We confirmed this prediction by performing FFPE-CUTAC on a set of 30 human meningiomas and showing that RNAPII at the 64 RC Histone genes, which comprise only 1/100,000^th^ of the human genome, successfully estimated WHO grade and accurately predicted rapid recurrence, which corresponds to elevated expression of proliferation genes (*33*). In sharp contrast, predictions based on RNAPII at Ribosomal Protein genes or mitochondrial DNA abundance failed. Although meningiomas are not invasive, elevated RNAPII at RC histone genes was also observed in invasive breast tumors. The ability of RNAPII FFPE-CUTAC at only the 64 human RC genes to predict aggressiveness in a common intracranial tumor and in multiple breast cancer subtypes implies that a rapid PCR assay for RC histone gene RNAPII or transcription (*39*) may become an inexpensive general cancer diagnostic tool, revolutionizing precision oncology.

We also found that levels of RNAPII at the 64 RC histone genes correlated with total aneuploidies, which are present in nearly all meningioma patient samples (*33*), consistent with the occurrence of ∼90% whole-arm imbalances in pan-cancer TCGA data (*35*). Intriguingly, whole-arm losses were observed to be in excess of gains for 38 of the 39 autosomal arms for both meningiomas and breast tumors representing multiple subtypes. Analysis of TCGA data uncovered no evidence that mitotic segregation errors could account for biases of losses over gains. To explain how over-production of histones might account for this striking whole-arm loss bias in patient tumors, we propose that excess H3 histones compete with CENP-A histones at S-phase for nucleosome assembly at centromeres (*40, 41*) (**Fig. 6D**). Production of S-phase histones is tightly regulated (*42–44*) and overproduction in cancer and displacement of CENP-A nucleosomes are known to result in the generation of DNA–RNA hybrids, likely due to transcription–replication conflicts causing delayed DNA replication, centromere breakage and loss of whole chromosome arms (*45, 46*). Thus, RNAPII excess over histone genes at S-phase provides a mechanistic basis for understanding not only how cancer cells can proliferate faster than their neighbors but also how the same process might generate centromeric breaks that result in whole-arm imbalances that drive most cancers.

## Acknowledgements

We thank Christine Codomo and Terri Bryson for technical assistance, the Fred Hutch Genomics Shared Resource for sequencing and data processing and the Fred Hutch Experimental Histopathology Shared Resource for FFPE processing and analysis, Oregon Health Sciences University Knight Biolibrary for breast tumor FFPEs, and Ryan Corces for advice on ATAC-seq data availability. This work was supported by the Howard Hughes Medical Institute (S.H.), National Institutes of Health grant HG012797 (Y.Z.), and grant # T32CA009515 from the National Cancer Institute (R.M.P.).

## Author Contributions

Conceptualization: SH, YZ, KA

Methodology: SH, YZ, JEG, JGH, HNT

Investigation: SH, YZ, RMP, YX, JEG, JGH, ZRR, FS, HNT, KA

Funding acquisition: SH Supervision: SH, ECH, SK, KA

Writing – original draft: SH

Writing – review & editing: SH, YZ, JGH, SK, ECH, KA

## Data and Materials Availability

The sequencing data generated in this study have been deposited in the NCBI GEO database under accession code GSE261351. The raw sequencing data generated from the University of Washington samples are not made available due to data privacy laws. Processed FFPE-CUTAC data for these samples have been deposited on Zenodo and can be accessed with the identifier https://doi.org/10.5281/zenodo.13138686. Custom scripts used in this study are available from GitHub: https://github.com/Henikoff/FFPE.

## List of Supplementary Materials Materials and Methods

### Ethical statement

This research was approved by the Fred Hutch Institutional Animal Care and Use Committee (Protocol # 50842) and complies with all required ethical regulations.

### Mouse tumor and normal tissues and FFPEs

Ntva;cdkn2a-/-mice were injected intracranially with DF1 cells infected with and producing RCAS vectors encoding either PDGFB (*16*), ZFTA-RELA (*14*), or YAP1-FAM118b (*15*) as has been described (*47*). Upon weaning (∼P21), mice were housed with same-sex littermates, with no more than 5 per cage and given access to food/water *ad libitum*. When the mice became lethargic and showed poor grooming, they were euthanized and their brains removed and fixed at least 48 hours in Neutral Buffered Formalin. All animal experiments were approved by and conducted in accordance with the Institutional Animal Care and Use Committee of Fred Hutchinson Cancer Center (Protocol #50842: Tva-derived transgenic mouse model for studying brain tumors). Tumorous and normal brains were sliced into five pieces and processed overnight in a tissue processor, mounted in a paraffin block and 10-micron sections were placed on slides. Mouse tissue (including normal and tumor bearing brains) was removed, fixed in 10% neutral-buffered formalin for a minimum of 24 hours and embedded into paraffin blocks. 10-µm serial sections were cut from formalin-fixed paraffin-embedded specimens and mounted on slides.

### Human FFPE slides

The following pairs of human tumor and adjacent normal 5-µm tissue sections on slides from single FFPE blocks were purchased from Biochain, Inc: Breast Normal/Tumor cat. no. T8235086PP/PT; Colon Normal/Tumor cat. no. T8235090PP/PT; Kidney Normal/ Tumor cat. no. T8235142PP/PT; Liver Normal/Tumor cat. no. T8235149PP/PT; Lung Normal/Tumor cat.

no. T8235152PP/PT; Rectum Normal/Tumor cat. no. T8235206PP/PT; Stomach Normal/Tumor cat. no. T8235248PP/PT. Human primary liver tumor and normal samples were harvested from cases undergoing surgical resection at the University of Washington under the Institutional Review Board approved protocol and then subsequently deidentified.

A set of 31 human meningioma 4-µm tissue sections on slides from the same FFPE blocks used for both frozen-RNA-seq and FFPE-RNA-seq were obtained from the University of Washington as part of a larger study (*33*). An additional set of 15 human breast tumor 10-µm FFPE tissue sections (dates of collection 1999-2012) on slides were obtained from Oregon Health Sciences University Knight BioLibrary. Single slides were subjected to on-slide FFPE-CUTAC and scraped into multiple tubes for PCR to approximately equalize tissue amount between samples. At least one replicate sample from each patient sample provided sufficient data for analysis.

### Antibodies

Primary antibodies: RNAPII-Ser5p: Cell Signaling Technologies cat. no. 13523, lot 3; RNAPII-Ser2p: Cell Signaling Technologies cat. no. 13499; H3K27ac: Abcam cat. no. ab4729, lot no. 1033973. Secondary antibody: Guinea pig α-rabbit antibody (Antibodies online cat. no. ABIN101961, lot 46671).

### On-slide FFPE-CUTAC

On-slide FFPE-CUTAC was performed as described (*11*) with modifications. The combination of several days in 4% formaldehyde in preparing FFPEs makes the vast majority of the genome impermeable and therefore not recovered. However, the RNAPII-Ser-5phosphate epitope, present the 52 tandem copies of the heptapeptide YSPTSPS (364 amino acids without a single lysine) is available, and to a first approximation only over active promoters, enhancers and very highly expressed genes such as the RC histone genes. Briefly, FFPE slides were placed in 800 mM Tris-HCl pH8.0 in a slide holder and incubated at 85-90°C for 1-14 hours, whereupon the paraffin melted and floated off the slide. Slides were cooled to room temperature and transferred to 20mM HEPES pH 7.5,150mM NaCl. Slides were drained and excess liquid wicked off using a Kimwipe tissue. The sections were immediately covered with 20-60 µL primary antibody in Triton-Wash buffer (20mM HEPES pH 7.5,150mM NaCl, 2mM spermidine and Roche complete EDTA-free protease inhibitor) added dropwise. Plastic film was laid on top to cover and slides were incubated ≥2 hr incubation at room temperature (or overnight at ∼8°C) in a moist chamber. The plastic film was peeled back, and the slide was rinsed once or twice by pipetting 1 mL Tri-ton-Wash buffer on the surface, draining at an angle. This incubation/wash cycle was repeated for the guinea pig anti-rabbit secondary antibody (Antibodies On-line cat. no. ABIN101961) and for pAG-Tn5 preloaded with mosaic end adapters (Epicypher cat. no. 15-1117 1:20), followed by a Triton-Wash rinse and transfer of the slide to 10 mM TAPS pH 8.5. Tagmentation was performed in 5mM MgCl_2_, 10mM TAPS pH 8.5, 20% (v/v) N,N-dimethylformamide in a moist chamber and incubated at 55°C for 1 hr. Following tagmentation, slides were dipped in 10 mM TAPS pH 8.5, drained and excess liquid wicked off. Individual sections were covered with 2 µL 10% Thermolabile Proteinase K (TL ProtK) in 1% SDS using a pipette tip to loosen the tissue. Tissue was transferred to a thin-wall PCR tube containing 2 µL TL ProK using a watchmaker’s forceps, followed by 1 µL TL ProtK and transfer to the PCR tube. Tubes were incubated at 37°C for 30 min and 58°C for 30 min before PCR as described above.

### FFPE-CUTAC for curls

Curls were transferred to a 1.7 mL low-bind tube (Axygen cat. no. MCT-175-C), which tightly fits a blue pestle (Fisher cat. on. 12-141-364). Mineral oil (200 µl) was added and the tube was placed in a 85-90°C water bath for up to 5 min to melt the paraffin. The suspension was then homogenized ∼10-20 sec with a pestle attached to a pestle motor (DWK Life Sciences cat no. 749540-0000). Warm cross-link reversal buffer (200 µl 800 mM Tris-HCl pH8.0) was added followed by addition of 6 µl of 1:10 Biomag amine paramagnetic beads (48 mg/ml, Polysciences cat. no. 86001-10). Homogenization was repeated, and 800 µl warm cross-link reversal buffer was added. Tubes were incubated at 85-90°C for 1-14 hours, vortexed, centrifuged briefly and the mineral oil was removed from the top without disturbing the surface. A 500 µl volume of mineral oil was added, mixed by inversion, centrifuged and the mineral oil removed leaving a thin oil layer. A 2.4 µl volume of agarose glutathione paramagnetic beads (Fisher cat. no. 88822) was added below the surface and mixed by inversion on a Rotator. Tubes were centrifuged briefly, placed on a strong magnet (Miltenyi Macsimag separator, cat. no. 130-092-168), and the supernatant removed and discarded, and the bead-bound homogenate was resuspended in up to 1 mL Triton-wash buffer (20 mM HEPES pH 7.5, 150 mM NaCl, 0.5 mM spermidine, 0.2mM EDTA, 0.05% Tri-ton-X100 and Roche EDTA-free protease inhibitor) and divided into PCR tubes for antibody addition. Other steps through to library preparation and purification followed the standard FFPE-CUTAC protocol (*11*). De-tailed step-by-step protocols for both slides and curls are available on Protocols.io: https://www.protocols.io/edit/cutac-for-ffpes-c5huy36w.

### DNA sequencing and data processing

The size distributions and molar concentration of libraries were determined using an Agilent 4200 TapeStation. Barcoded CUTAC libraries were pooled at equal volumes within groups or at approximately equimolar concentration for sequencing. Paired-end 50×50 bp sequencing on the Illumina NextSeq 2000 platform was performed by the Fred Hutchinson Cancer Center Genomics Shared Resources.

### Data analysis

#### Preparation of the cCREs

We obtained the mm10 and hg38 versions of the Candidate *cis*-Regulatory Elements (cCREs) by EN-CODE (https://screen.encodeproject.org/) from UCSC (*48*). For mouse mm10 we used all 343,731 entries. Because our sequencing data was aligned to hg19, we used UCSC’s liftOver tool to re-position the hg38 cCREs, resulting in 924,834 entries. We noticed that many human cCREs were in repeated regions of the genome, so we intersected the hg19 cCRE file with UCSC’s RepeatMasked regions using bedtools 2.30.0 (*49*) “intersect - v” command to make a file of 464,749 cCREs not in repeated regions.

#### Preparation of histone regions

For mm10 we used these regions:

|  |  |  |  |
| --- | --- | --- | --- |
| chr13 | 21715711 | 21837530 | H2bc13-H4bc2 |
| chr13 | 22035122 | 22043658 | H2ac12-H2bc11 |
| chr13 | 23531044 | 23622558 | H4c8-H1f4 |
| chr13 | 23683473 | 23764412 | H2ac6-H1f1 |

For hg19 we used the 64 annotated regions within the two histone clusters (**Table S2**):

|  |  |  |  |
| --- | --- | --- | --- |
| chr1 | 149783434 | 149859466 | Minor |
| chr6 | 26017260 | 26285727 | Major |

#### FFPE CUTAC samples alignment of PE50 Illumina sequencing

1. We used cutadapt 2.9 (*50*) with parameters *‘-j 8 --nextseq-trim 20 -m 20 -a AGATCGGAAGAGCA-CACGTCTGAACTCCAGTCA -AAGATCGGAAGAG-CGTCGTGTAGGGAAAGAGTGT -Z*’ to trim adapters from 50bp paired-end reads FASTQ files.
2. We used Bowtie2 2.4.4 (*51*) with options *‘--very-sen-sitive-local --soft-clipped-unmapped-tlen –dovetail --no-mixed --no-discordant -q --phred33 -I 10 -X 1000*’ to map the paired-end 50bp reads to the mm10 Mus musculus or hg19 Homo sapiens reference sequences obtained from UCSC.
3. We used samtools 1.14 (*52*) ‘*view*’ to extract prop-erly paired reads from the mm10 alignments into BED files of mapped fragments.
4. We computed the fraction of fragments mapped to chrM.
5. We used bedtools 2.30.0 ‘*genomecov*’ to make a normalized count track which is the fraction of counts at each base pair scaled by the size of the reference sequence so that if the counts were uniformly distributed across the reference sequence there would be one at each position.
6. We ran Picard 2.18.29 ‘*MarkDuplicates*’ program (http://broadinstitute.github.io/picard/) on the SAM out-put files from bowtie2.

#### RNA-seq public samples re-alignment

We re-analyzed 1298 meningioma patient samples, which included raw FASTQ files and clinical outcomes, obtained from Thirimanne et al. (2024). For alignment, we used the reference genome hg19 from GENCODE (v19). The RNA-seq data was aligned using STAR version 2.7.11 with the following parameters for per-sample 2-pass mapping and we utilized the unstranded RNA-seq counts as the raw count per gene per sample:

--outSAMtype BAM Unsorted
--outSAMattributes NH HI NM MD AS nM
--twopassMode Basic
--twopass1readsN −1
--quantMode TranscriptomeSAM GeneCounts
--readFilesCommand zcat

#### Preparation of aligned samples

1. For mouse on-slide samples, we merged tumor replicates within the experiment. The raw count was normalized with respect to the sequencing depth and genome coverage prior to generating the genome browser track visualization and conducting the hypertranscription analysis.
2. For BioChain human on-slide tumor and normal samples, we merged mapped fragments from different experiments for each tumor or normal. We then equalized the numbers of fragments for tumor and normal pairs by downsampling the larger set of the two using the UNIX ‘*shuf*’ command. This ensured that each matching tu-mor-normal sample comparison was based on the equal total number of reads.
3. Meningioma FFPE-CUTAC samples, with RNAPII enrichment counts taken at 500bp genomic binning, cCRE regions, and genes respectively, were normalized by the term fre-quency-inverse document frequency (TF-IDF) normalization method (*53*) to remove the sequencing depth biases.

#### Peak calling

We ran SEACR 1.3 (*18*) with parameters ‘*norm re-laxed*’ on tumor samples with the normal sample from each tumor and normal pair as the control. For comparison, we also called peaks after reversing the roles of tumor and normal.

#### Preparation of and Tumor-Normal file per cCRE region

We used the bedtools ‘*intersect*’ and ‘*groupby*’ com-mands to sum normalized counts within the cCRE region boundaries. The resulting files have one row per cCRE region and one column per sample and are suitable for submission to the Degust server (https://de-gust.erc.monash.edu/) using the ‘*Voom/Limmà* option (-log_10_FDR versus log_2_FoldChange). We computed Tu-mor-Normal pairs from the cCRE region files and sorted them by largest differences in absolute value (**Table S1**).

#### Curve-fitting

We partitioned the genome into 1 kb tiles and merged replicates, then downsampled to equalize library sizes between tumor and normal samples from each patient and added up normalized counts within each tile. For each tumor and normal patient sample, we fit the normalized counts across tiles using a Local Polynomial Regression (LOESS) model as implemented in the ‘*stats*’ package of the R programming language, setting the degree of smoothing to 0.2 specified by the ‘*span*’ parameter of ‘*loess*’ function.

#### UMAPs

To ensure the quality of samples for downstream analysis, we excluded BioChain tumor or normal samples with fewer than 100,000 read counts or less than 10,000,000 bp of total fragment length. We utilized 500bp genomic binning, cCRE regions, or gene regions as the genomic features, respectively. Specifically, each gene feature was defined as the region spanning from the 3’-most transcript end through the 5’-most end, stopping when either end of the next gene or LINE element s reached. For histone gene features related analysis, we took the 64 corresponding histone gene regions from the all-gene regions defined as above. We calculated the raw sequencing read count overlapping each type of feature regions using the ‘*getCounts*’ function from the chromVAR R package. Processing the feature regions by samples count matrix, we initially applied the TF-IDF normalization method (*53*). This method first normalizes read counts across samples to correct for differences in total read depth and then adjusts across cCRE regions, assigning higher values to rarer regions. TF-IDF normalization was implement-ed using the ‘*RunTFIDF*’ function from the Signac R package. This step was followed by the selection of top features using ‘*FindTopFeatures*’ from the Signac package and data scaling performed by ‘*ScaleData*’ from the Seurat package. Subsequently, we conducted principal component analysis (PCA) on the scaled data and then used the top 50 principal components to generate a UMAP representation, providing a refined visualization of the relationship across samples.

#### Meningioma WHO grade prediction

All 64 gene features are used to generate the UMAP to demonstrate the separation power of histone gene between tumor and normal samples. The average distance to all the normal samples are calculated on 50 dimensional latent embeddings using Euclidean distance for each tumor sample. The FFPE-CUTAC meningioma samples were then ranked based on the distance to normal samples, with longer distance ranked higher and shorter distance ranked lower. This ranking was subsequently correlated with the corresponding WHO grade of meningioma patients.

#### FFPE-CUTAC and RNA-seq meningioma samples integration

We leveraged the canonical correlation analysis strategy (*53*) to integrate the FFPE-CUTAC samples with the public RNA-seq meningioma data. First, we quantified the FFPE-CUTAC RNAPII enrichment signals across genes to obtain a gene-by-sample count matrix in the same format as the RNA-seq processed data. Both FFPE-CUTAC and RNA-seq data were normalized by log-transformation of counts per million (CPM), center-scaled, and subjected to principal component analysis (PCA) on the top variable features. For the integration, we retained almost all the genes using the top 5,7000 features and the top 250 PCs for dimen-sion reduction. The ‘FindIntegrationAnchors’ and ‘In-tegrateDatà functions from the Seurat v5.1 R package were used to find the anchor samples between FFPE-CUTAC and RNA-seq, and to integrate two modalities. We set ‘k.anchor = 5’ to find the top 5 matching samples in the other modality. The integrated UMAPs were generated using 250 PCs and 15 nearest neighbors, with the remaining parameters set to default values.

#### Meningioma recurrence prediction

We defined the overall RNAPII signals across histone genes by taking the median of the TF-IDF normalized counts of 64 histone genes. The density distribution across 30 meningioma patients provided the general pattern of histone signals (**fig. S11A**). Considering that meningioma is mostly benign, we set the threshold at 4.4 to call a high histone signal at the right tail of the highest density peak. Five patients are hence deemed to have high RNAPII enrichment at histone genes for the prediction of faster recurrence. For a fair comparison, the top five patients were also used for the ribosomal protein genes and the fraction of Chromosome M (chrM) prediction (**Fig. 4B-D**).

We next investigated whether the high histone signal could predict recurrence outcomes. Due to the lack of clinical records for the 30 meningioma patients profiled by FFPE-CUTAC, we leveraged public frozen RNA-seq data of meningioma, which includes recurrence time and status information. These RNA-seq data have been successfully integrated and constructed into a shared nearest neighbor graph with our FFPE-CUTAC samples. The top 20 nearest neighbors were collected for each of the FFPE-CUTAC patient samples, and the Kaplan-Meier curves were generated for high and low histone signal groups, respectively. Log-rank test was used to compare the survival curves and determine if the recurrence rate in the high-histone-signal population was significantly faster than the low-histone-signal group. We implemented the same analysis on ribosomal protein genes and the fraction of chrM as negative controls and also explored the prediction performance using a series of thresholds to define the malignancy group.

### Aneuploidy prediction of meningioma recurrence

CaSpER was used to detect chromosome arm gain or loss for meningioma patients through the matching RNA-seq data. We followed the CaSpER tuto-rial (https://github.com/akdess/CaSpER?tab=read-me-ov-file) (*54*) demonstrated using the meningioma data from the Yale study (*55*). We used the same control samples, i.e., SRR3996005, SRR3996002, SRR3995998, SRR3996001, SRR3995989, SRR3995987, SRR3996004 and SRR3995995, from GEO: GSE85133. The same default parameter value, 0.75, was used when calling chromosome arm gain or loss using the function ‘extract-LargeScaleEvents’ from the CaSpER R package.

---

**Fig. S1.**
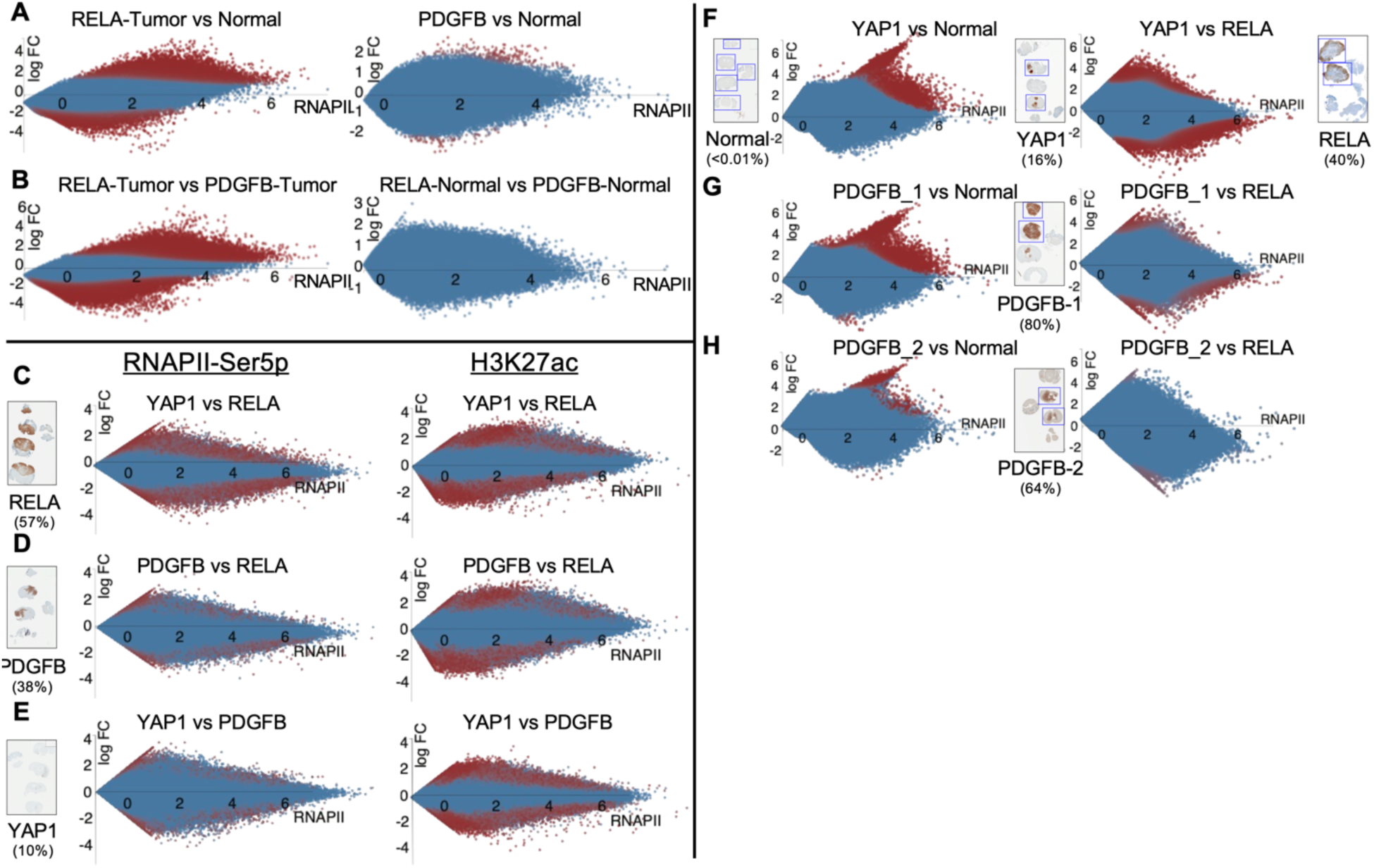
RNAPII-Ser5p FFPE-CUTAC shows stronger and more frequent changes in up-regulation than down-regulation of cCREs. The Voom/Limma option of the Degust server (https://degust.erc.monash.edu/) was applied to mouse cCRE RNAPII-Ser5p FFPE-CUTAC data from pooled replicates from 5 RELA and 4 PDGFB experiments. MA plots display *x* = log_10_(Tumor*Normal)/2 versus *y* = log_2_(Tumor/Normal) for normalized counts from the tumor and normal samples being compared, and red color indicates FDR < 0.05. Normalized counts are the fraction of counts at each base pair scaled by the size of the Mm10 reference sequence (2,818,974,548), so that if the counts are uniformly distributed across the reference sequence, there would be one at each position. (**A-B**) Both RELA and PDGFB tumor sections show higher counts than normal sections but significant RELA changes both up and down are far stronger than PDGFB changes, confirmed in a head-to-head comparison between tumors and normal sections. (**C-E**) Same as (A-B) except using either RNAPII-Ser5p or histone H3K27ac antibodies for FFPE-CUTAC and using entire 10-µm curls divided into 4-8 samples per curl for PCR and sequencing. For MA plots, data were merged from multiple experiments and equalized by downsampling to 10 million fragments, with 4 merged replicates per sample. DAP-stained slides for each paraffin block used, with the total fraction of tumor indicated in parentheses. (**F-H**) Voom/Limma was used to construct MA plots based on individual 10-µm sections from single slides corresponding to the boxed sections on slides DAP-stained for tumor-driver transgene expression. Numbers in parentheses are percentages of tumor cells based on the numbers of stained and unstained cells within the boxed sections. Relative to normal, more cCREs increase in RNAPII than decrease.

**Fig. S2.**
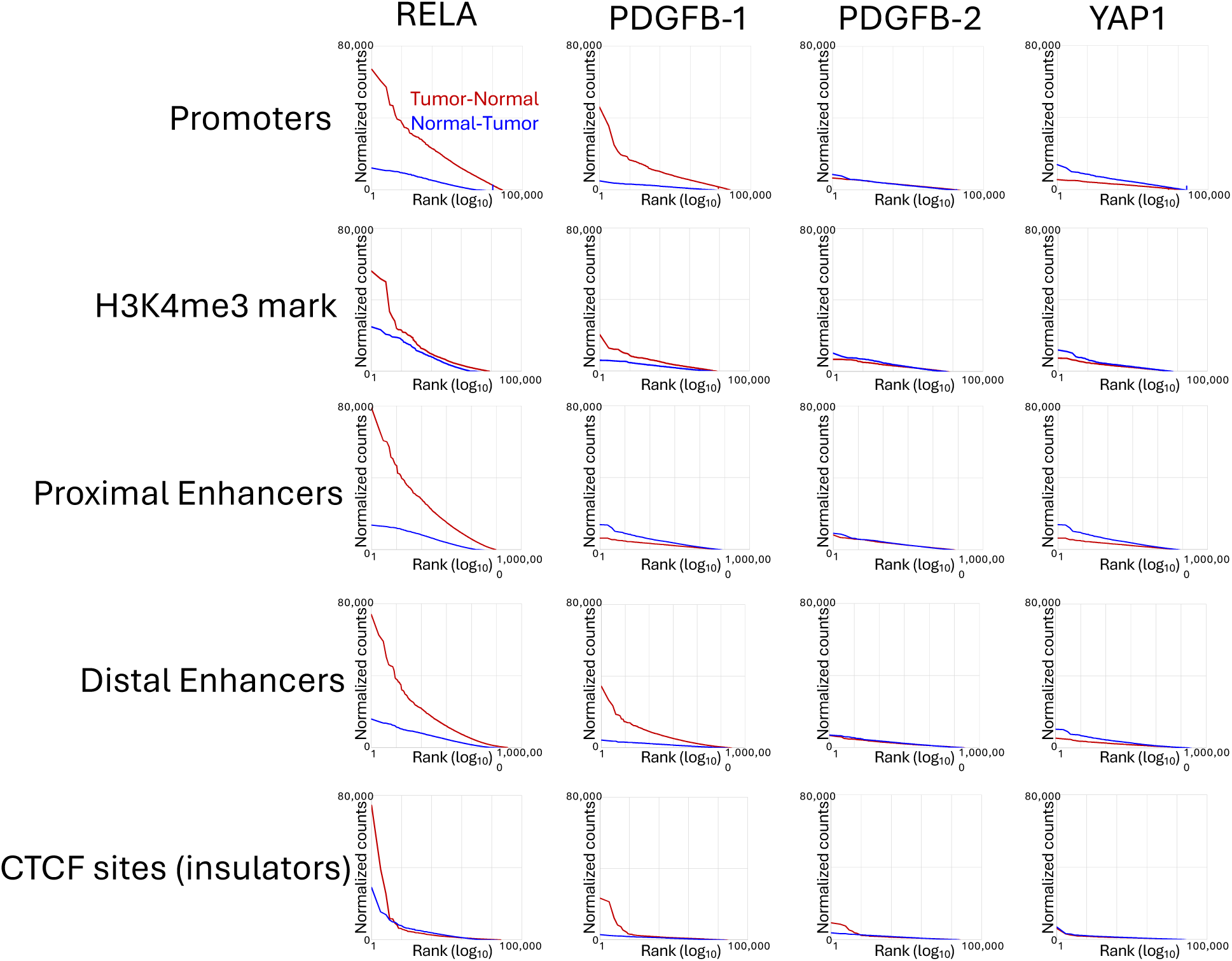
Hypertranscription mapped over the 343,731 ENCODE-annotated mouse cCREs categorized by regulatory element type. For each tumor and normal sample, we counted the number of mapped fragments spanning each base-pair in a cCRE scaled to the mouse genome and averaged the number of counts over that cCRE across tumor or normal samples. We then divided up the 343,731 cCREs into the five ENCODE-annotated categories: Promoters (24,114), H3K4me3-marked cCREs (10,538), Proximal Enhancers (108,474), Distal Enhancers (211,185) and CTCF cCREs (24,072) and rank-ordered based on tumor minus normal representing global upregulation (red), and conversely rank-ordered cCREs based on normal minus tumor representing global downregulation (blue). With such a large collection of loci, our a priori expectation is that the rank-ordered distribution of differences between tumor and normal will be approximately the same regardless of whether the differences are based on tumor minus normal or normal minus tumor. For clarity, we plotted rank-ordered differences on a log_10_ scale. Strong hypertranscription for RELA and PDGFB-1, weak hypertranscription for PDGFB-2, and little or no hypotranscription for YAP1 are seen for all classes, consistent with the T – N versus log_10_(T + N)/2 plots shown in Figure 1B-E.

**Fig. S3.**
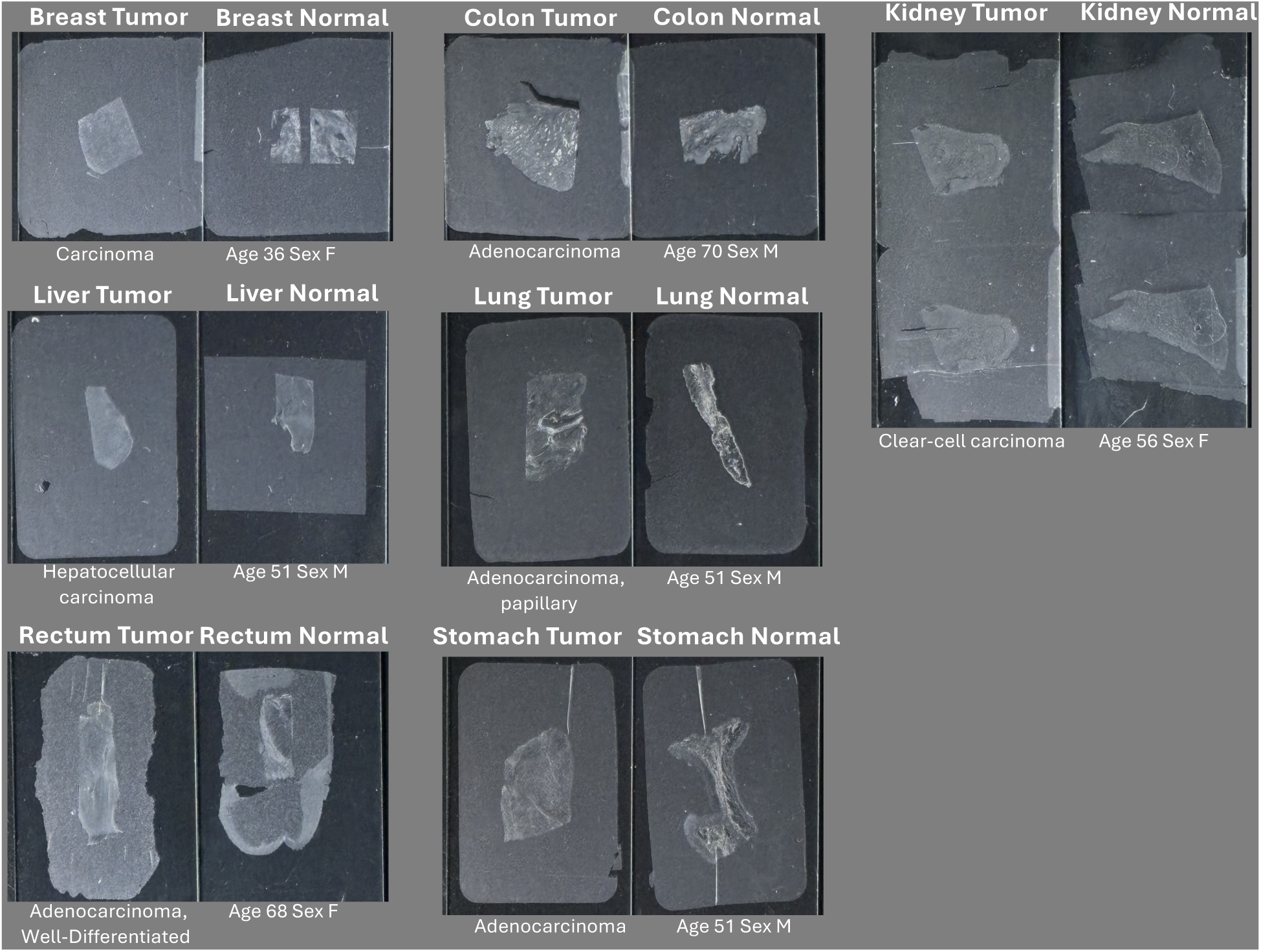
Photographs of 5-µm FFPE sections from human tumor and adjacent normal tissues. Pathology classification, age and sex were provided by the vendor (BioChain). Each image spans the width of a standard charged microscope slide, where the tissue is visible under the paraffin skin. On-slide RNAPII-Ser5p FFPE-CUTAC was applied to slides in parallel, using a total of four slides each for 100 separate samples in all to produce the data analyzed in this study.

**Fig. S4.**
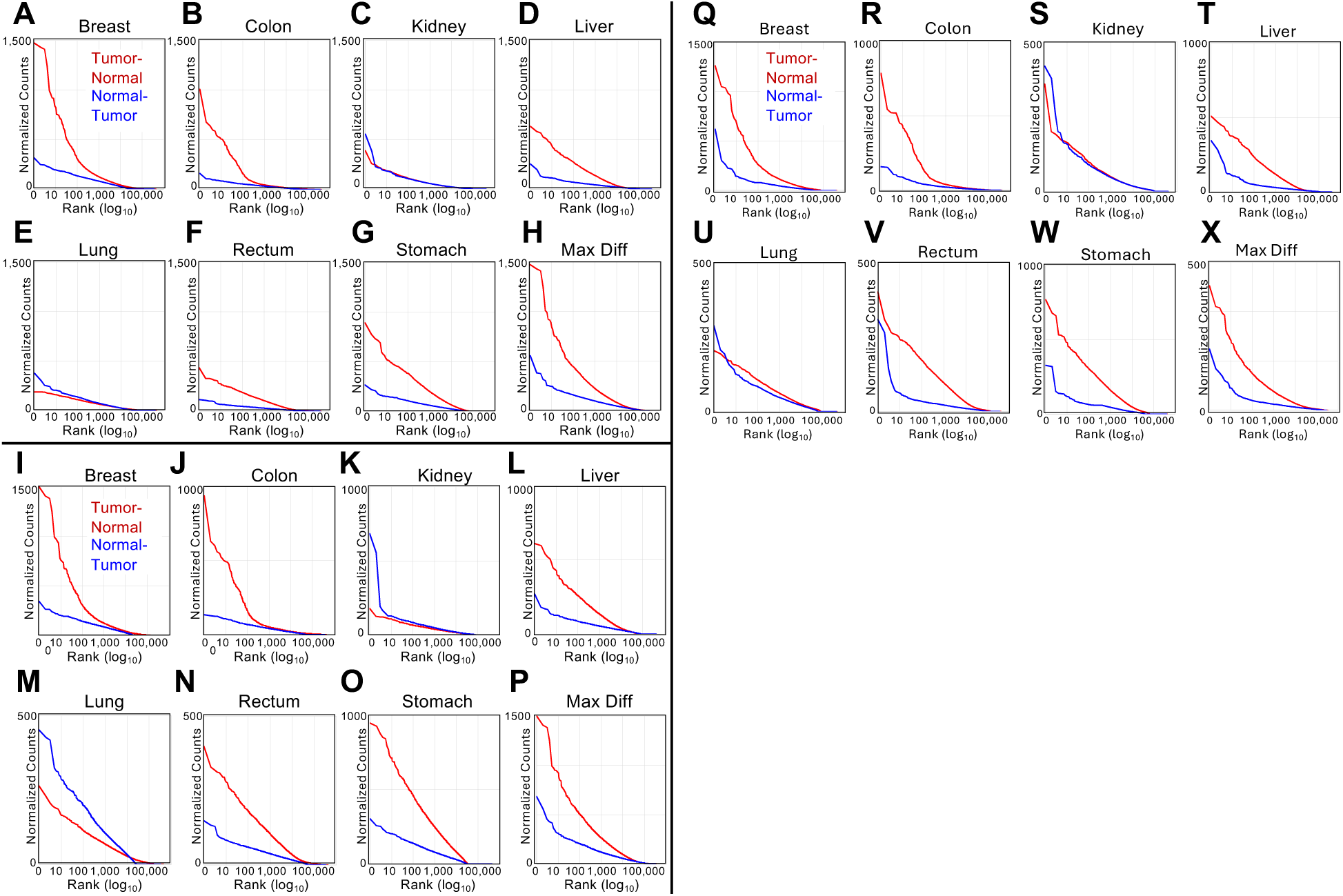
Hypertranscription in human Tumor-vs-Normal tissues. Same data as in the scatterplots of Figure 1F-M except line-plotted to facilitate comparisons. For each tumor and normal sample, we counted the number of mapped fragments spanning each base-pair in a cCRE scaled to the human genome and averaged the number of counts over that cCRE. We rank-ordered cCREs based on tumor minus normal representing global upregulation (red curves), and conversely rank-ordered cCREs based on normal minus tumor representing global downregulation (blue curves). With such a large collection of loci, our a priori expectation is that the rank-ordered distribution of differences between tumor and normal will be approximately the same regardless of whether the differences are based on tumor minus normal or normal minus tumor. For clarity, we plotted rank-ordered differences on a log_10_ scale. (**I-P**) Combined data from a single slide with duplicate removal. (**Q-X**) Combined data from four slides after removing duplicates and equalizing the number of fragments between tumor and normal sections. The number of unique fragments per sample in each Tumor/Normal pair is Breast: 1,125,608; Colon: 3,712,097; Kidney: 2,031,893; Liver: 2,983,411; Lung: 1,123,638; Rectum: 3,284,736; Stomach: 719,598.

**Fig. S5.**
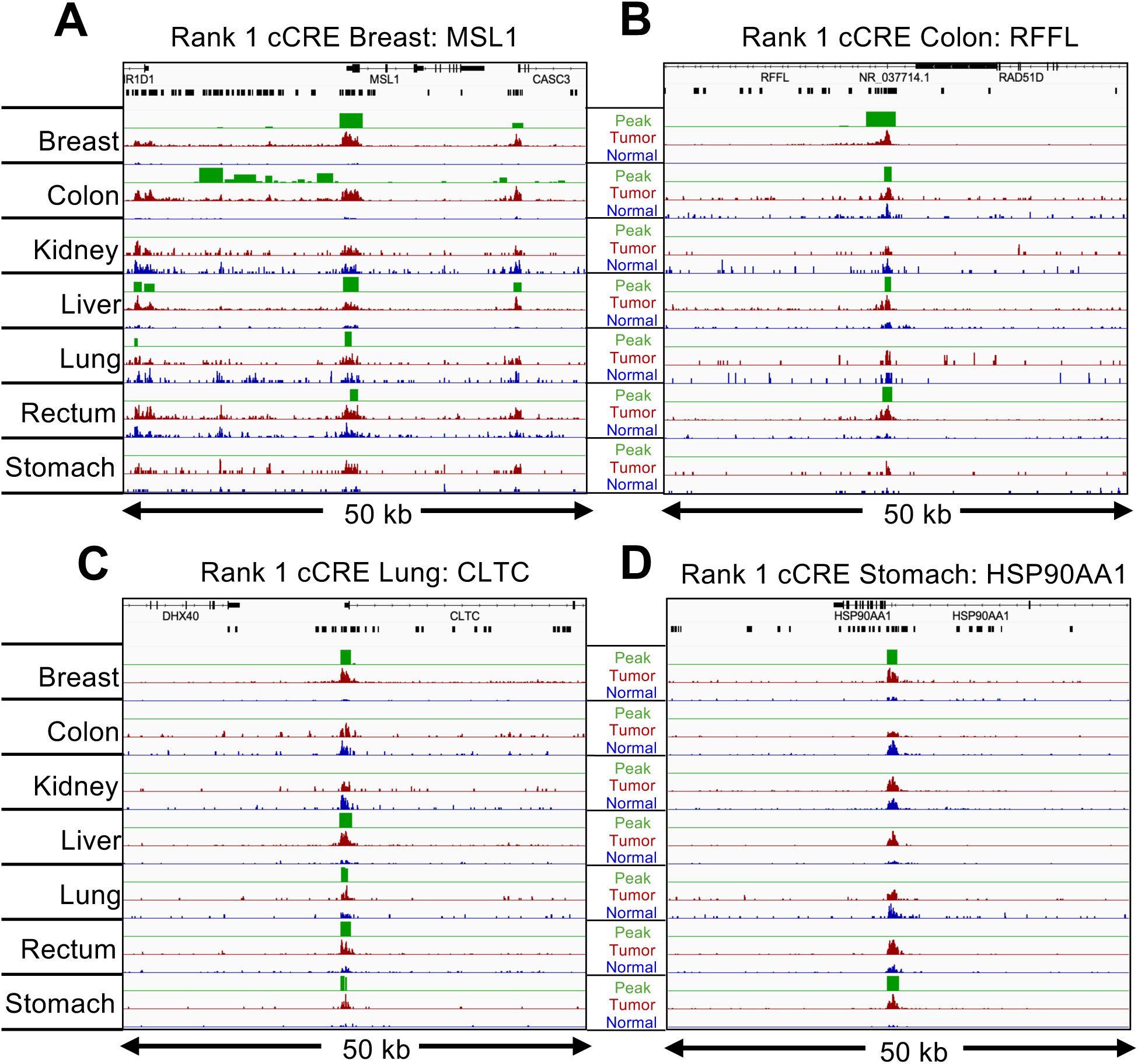
Genome browser tracks for the top-ranked CRE regions for Breast, Colon, Lung and Stomach. For each of the indicated tumors, tracks are shown for 50-kb regions around the #1-ranked cCRE based on Tumor (dark red) and Normal (blue) counts. Data tracks were group-autoscaled together for Tumor (red) and Normal (blue), where SEACR Tumor peak calls (green) use Normal as the negative control. Gene annotations and cCREs (black rectangles) are shown at the top. The #1-ranked cCREs intersected promoters in the Breast, Colon and Lung samples and intersected an intergenic enhancer in the HSP90AA1 gene in the Stomach sample.

**Fig. S6.**
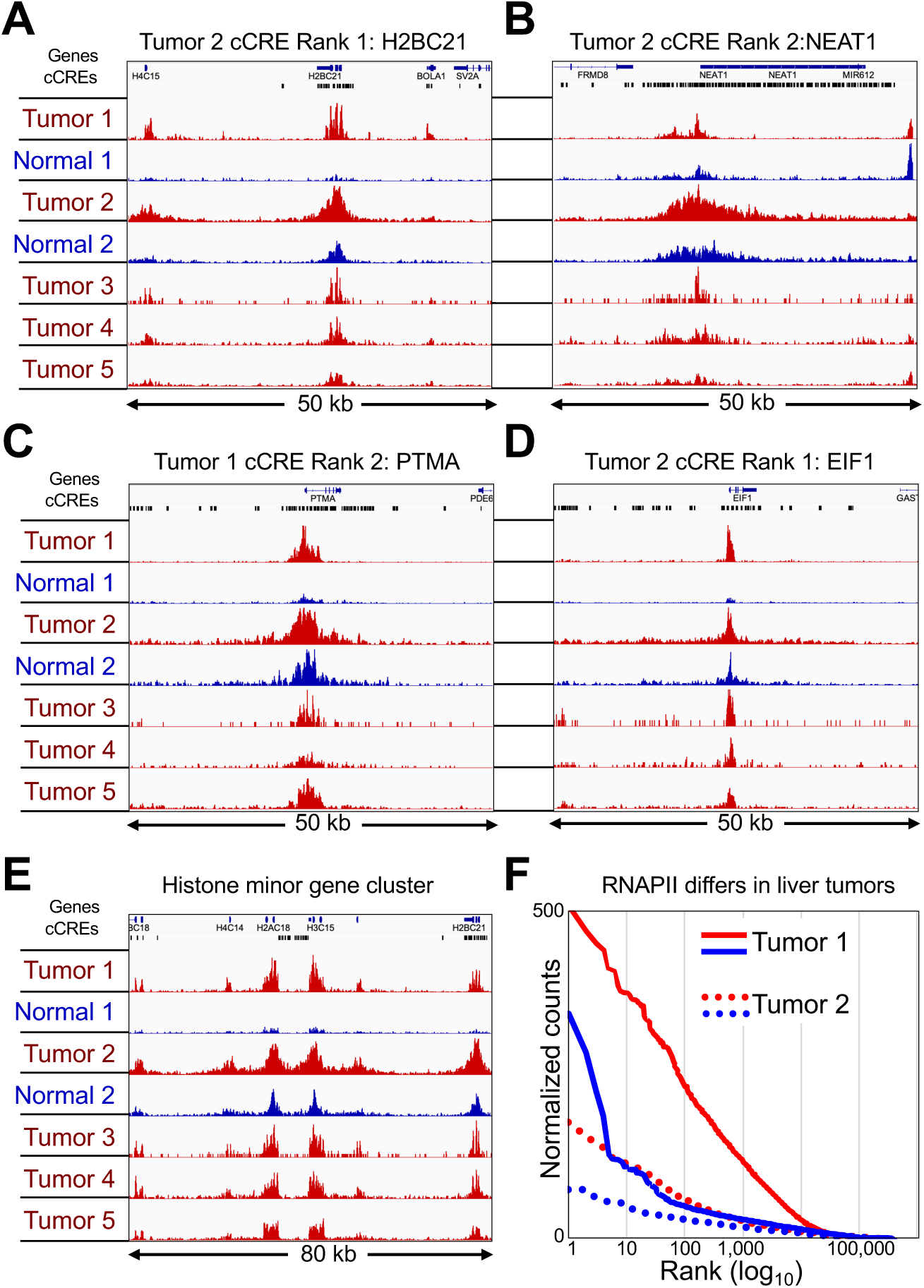
Hypertranscription differs between human liver tumors. (**A-D**) Top-ranked cCREs based on liver tumors 1 and 2 (red) and matched normal (blue) counts. Tumor/Normal tracks and Tumors 3-5 are group-autoscaled. (**E**) Same as (A), except for the minor histone gene cluster on Chromosome 1. (**F)** Levels of hypertranscription differ between different hepatocarcinomas (Tumor 1: solid lines, Tumor 2 dotted lines, where tumor is red and normal is blue) plotted as in Fig. S4.

**Fig. S7.**
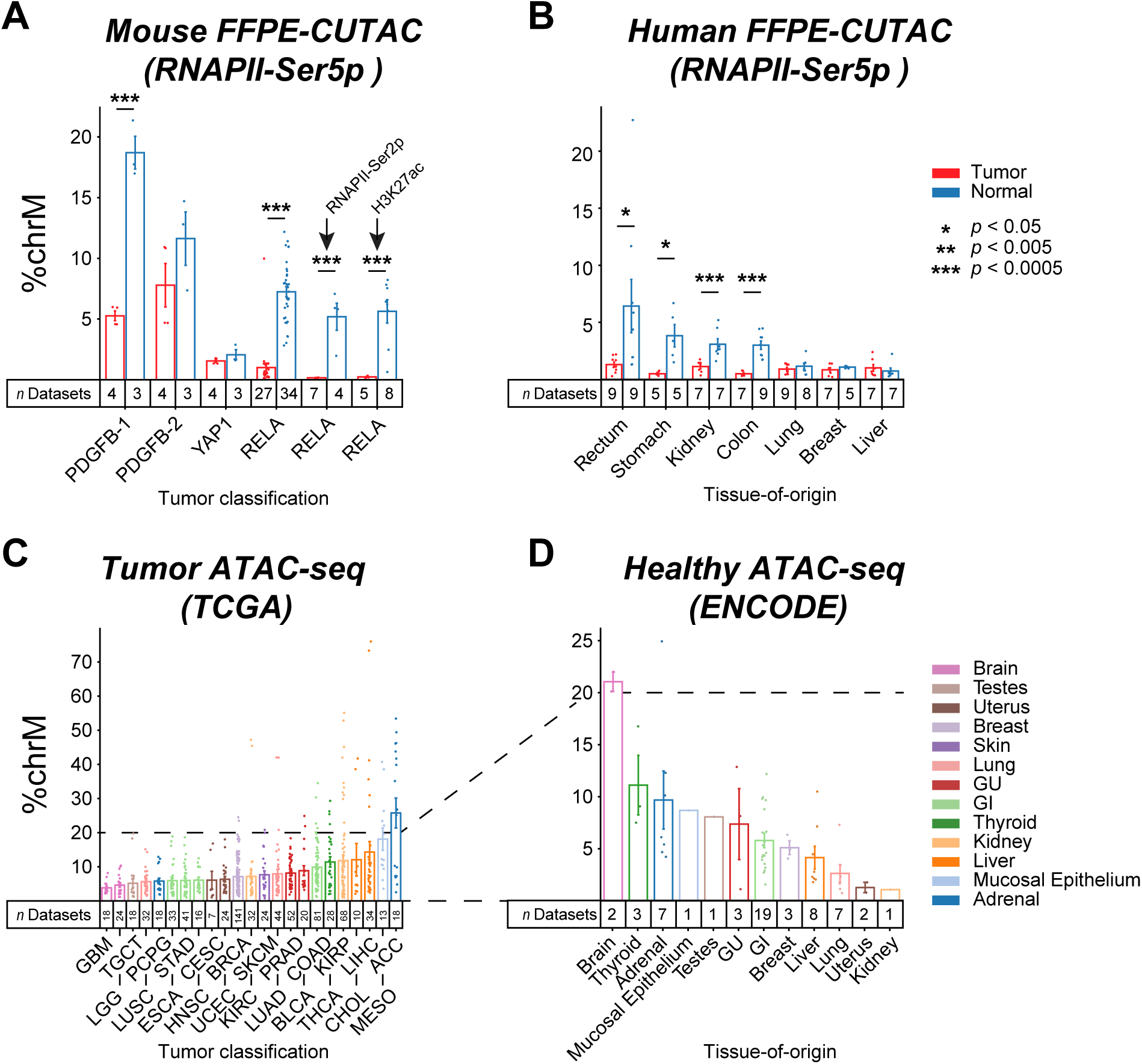
FFPE-CUTAC mitochondrial DNA signal is reduced in tumors. **A**) The percentage of normalized counts mapping to Chromosome M (chrM = mitochondrial DNA) was calculated for FFPE-CUTAC data from four mouse brain tumor paraffin blocks driven by PDGFB, YAP1 and RELA transgenes. An RNAPII-Ser5p antibody was used for the first four comparisons, and an RNAPII-Ser2p and histone H3K27ac antibodies were used respectively for the fifth and sixth comparisons. (**D**) Same as (C) for RNAPII-Ser5p FFPE-CUTAC data for the seven human Tumor/Normal pairs used in this study. (**E-F**) ATAC-seq count data from TCGA (tumor) and ENCODE (normal) shows variability in chrM percentages between tumors, consistent with our finding based on FFPE-CUTAC.

**Fig. S8.**
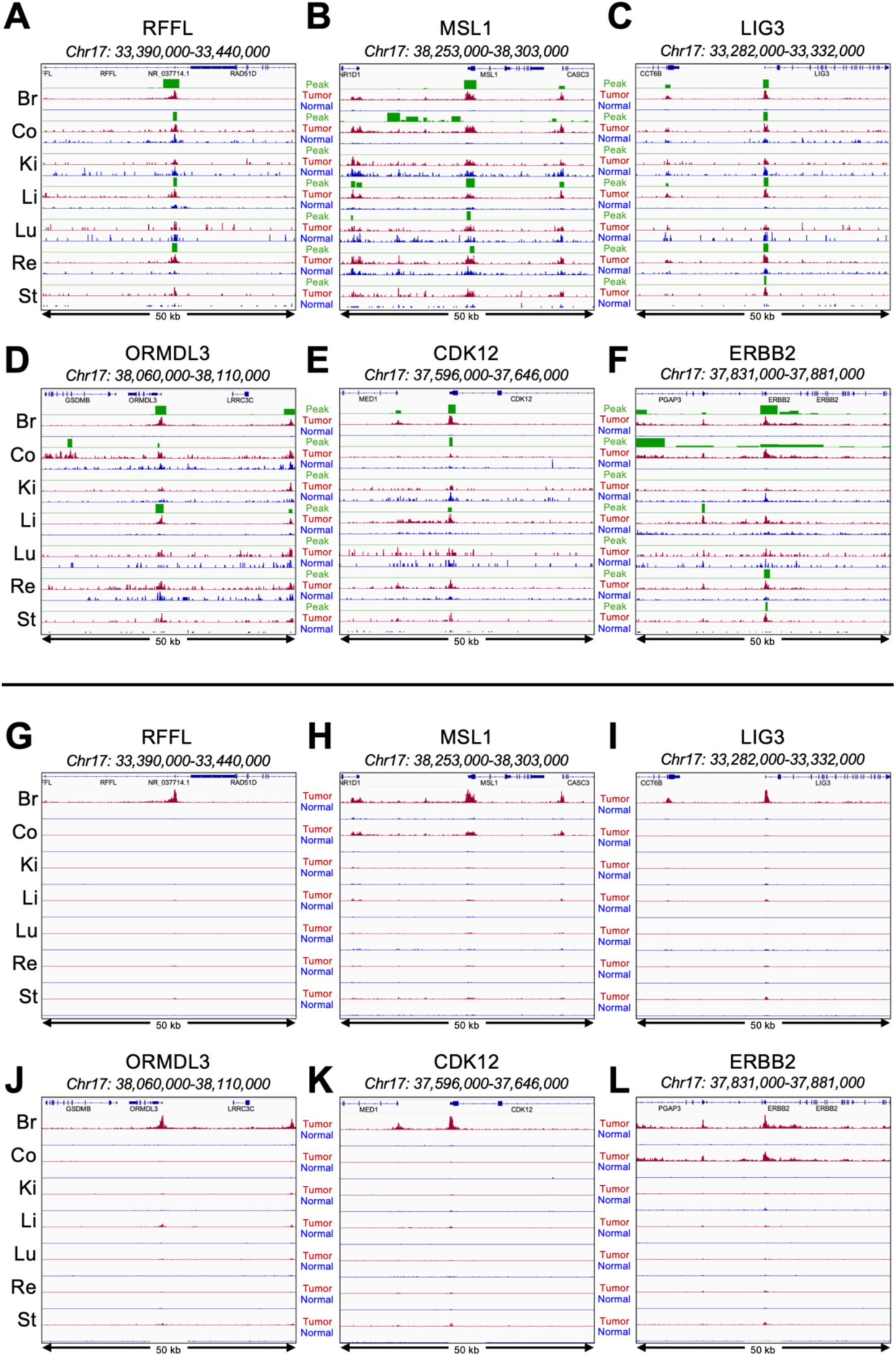
Focal hypertranscribed regulatory elements embedded in broad regions of hypertranscription on Chromosome 17q12-22. (**A-F**) The six most highly transcribed cCREs within the ∼5 Mb region of Chromosome 17q1.2-2.2 are displayed with each tumor (dark red) and normal (blue) pair scaled to one another so that peaks can be observed in all samples. SEACR peaks (green) are group-autoscaled in all panels. (**G-L**) Same as (A-F) except that all tumor-normal samples are group-autoscaled to the height of the tallest peak, where the disappearance of all the peaks except for those in Breast and for MSL1 and ERBB2 in Colon is evidence that peaks in these regions are strongly hypertranscribed in Breast and partially in Colon but not in any of the other tumors.

**Fig. S9.**
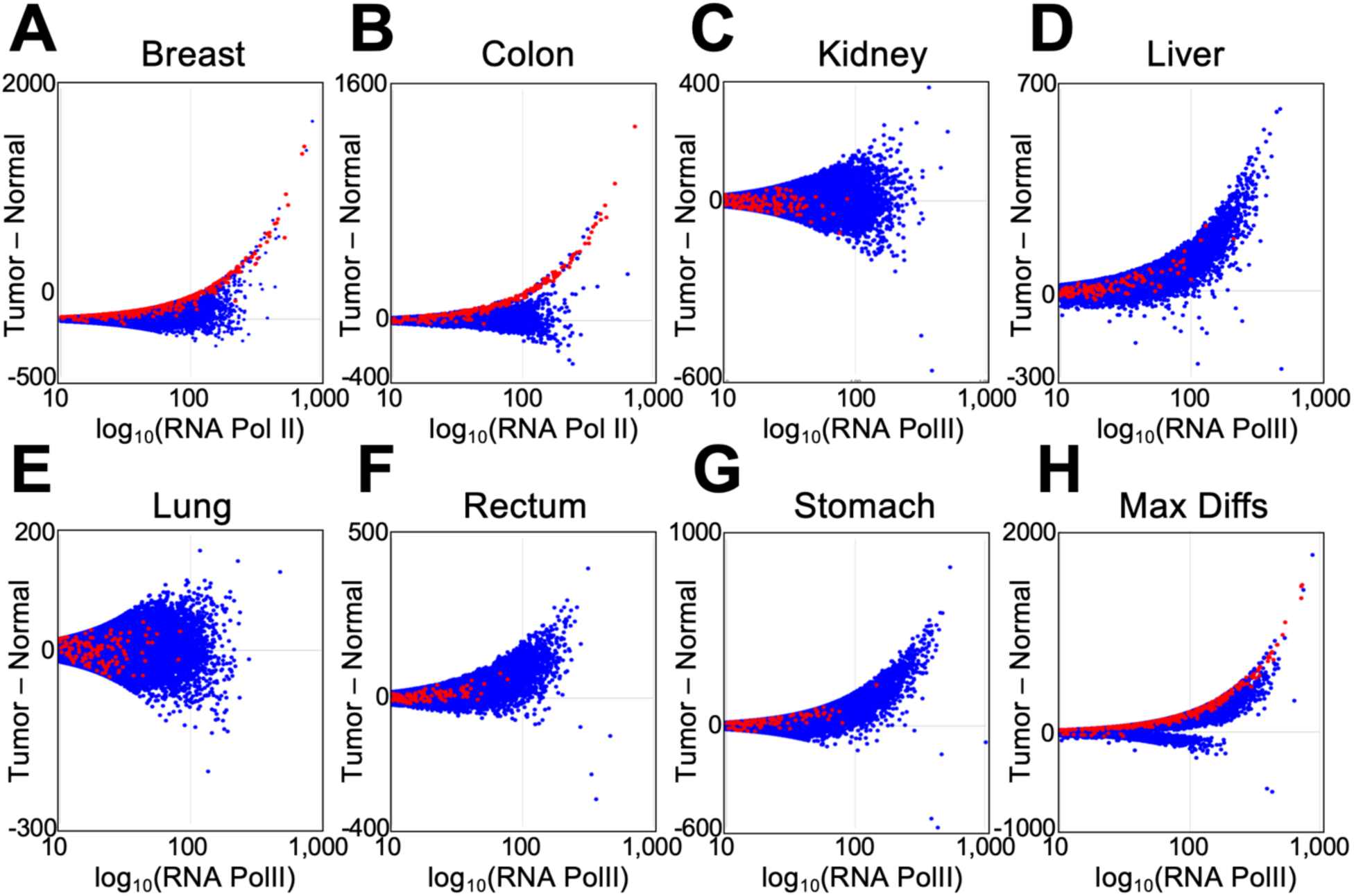
HER2 amplifications account for most of the hypertranscription signal in the Breast and Colon cancer samples. Panels A-H correspond to Figure 1F-M, where the superimposed red dots are cCREs within Chr17q12-21.

**Fig. S10.**
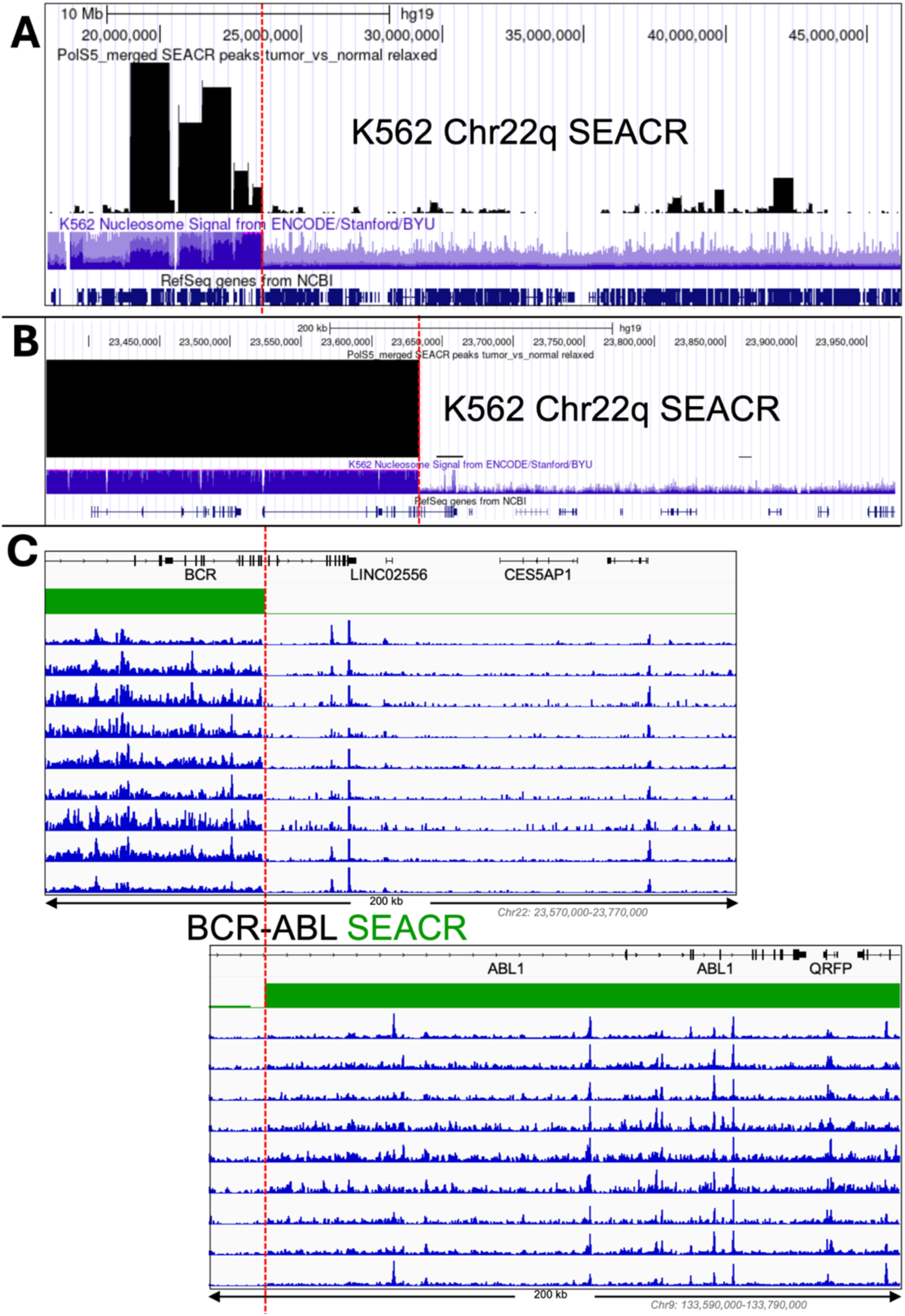
SEACR identifies and precisely maps amplifications in cancer. We called SEACR peaks on a merged set of RNAPII-Ser5p CUTAC datasets from K562 cells, which are annotated for amplifications in human genome build hg19. An amplified region (purple tracks) in K562 cells on Chr22q is shown as a UCSC browser track at (**A**) 25 Mb and (**B**) at 600 kb scales, together with SEACR broad peaks called on published K562 RNAPII-Ser5p CUTAC datasets (*20, 21*). Dotted red line indicates the location of the BCR-ABL t(9;22)(q34;q11) translocation breakpoint on Chr22q. (**C**) The precise correspondence to the annotated amplified regions is evident in the SEACR tracks and in 9 autoscaled tracks from 3 separate experiments, where the breakpoint in each gene (marked by the dotted red line) precisely corresponds to the change in signal amplitude, with abrupt increases within the part of each gene that is amplified, precisely mapped by SEACR.

**Fig. S11.**
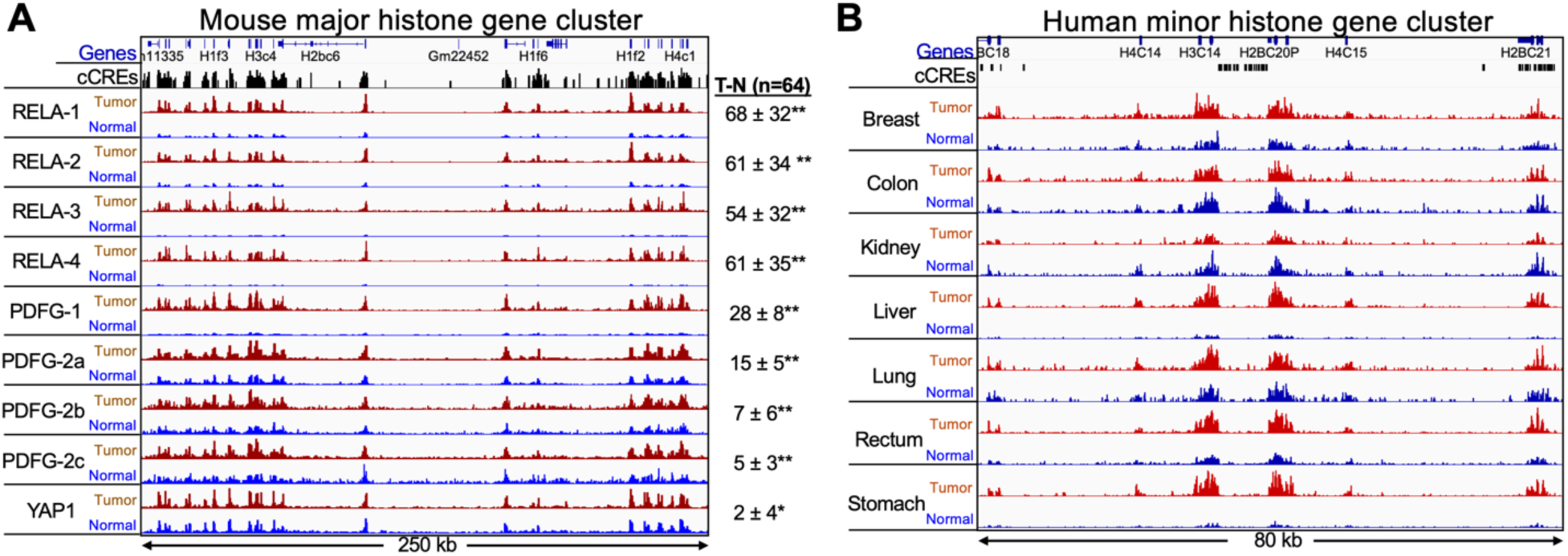
Hypertranscription at replication-coupled histone genes. (**A**) IGV tracks of the major histone gene cluster on Chromosome 13 from fusion-transgene-driven mouse brain tumors. Tumor and Normal 10-µm sections were from the same slide. Slides used for PDGFB-2a-c were from the same paraffin block but used in different experiments, and all others were from different paraffin blocks. Numbers at right were obtained by subtracting the sum of normalized counts in the normal sections from that in the tumor sections over all 64 annotated single-exon replication-coupled histone genes, where the Standard Deviation is shown. Paired *t*-test: * *p* < 0.001; ** *p* < 0.00001. (**B**) IGV tracks of the human minor histone gene cluster on Chromosome 1, where tracks are autoscaled for each Tumor (red) and Normal (blue). Tumor and Normal 5-µm sections were from a matched pair of slides taken from the same patient.

**Fig. S12.**
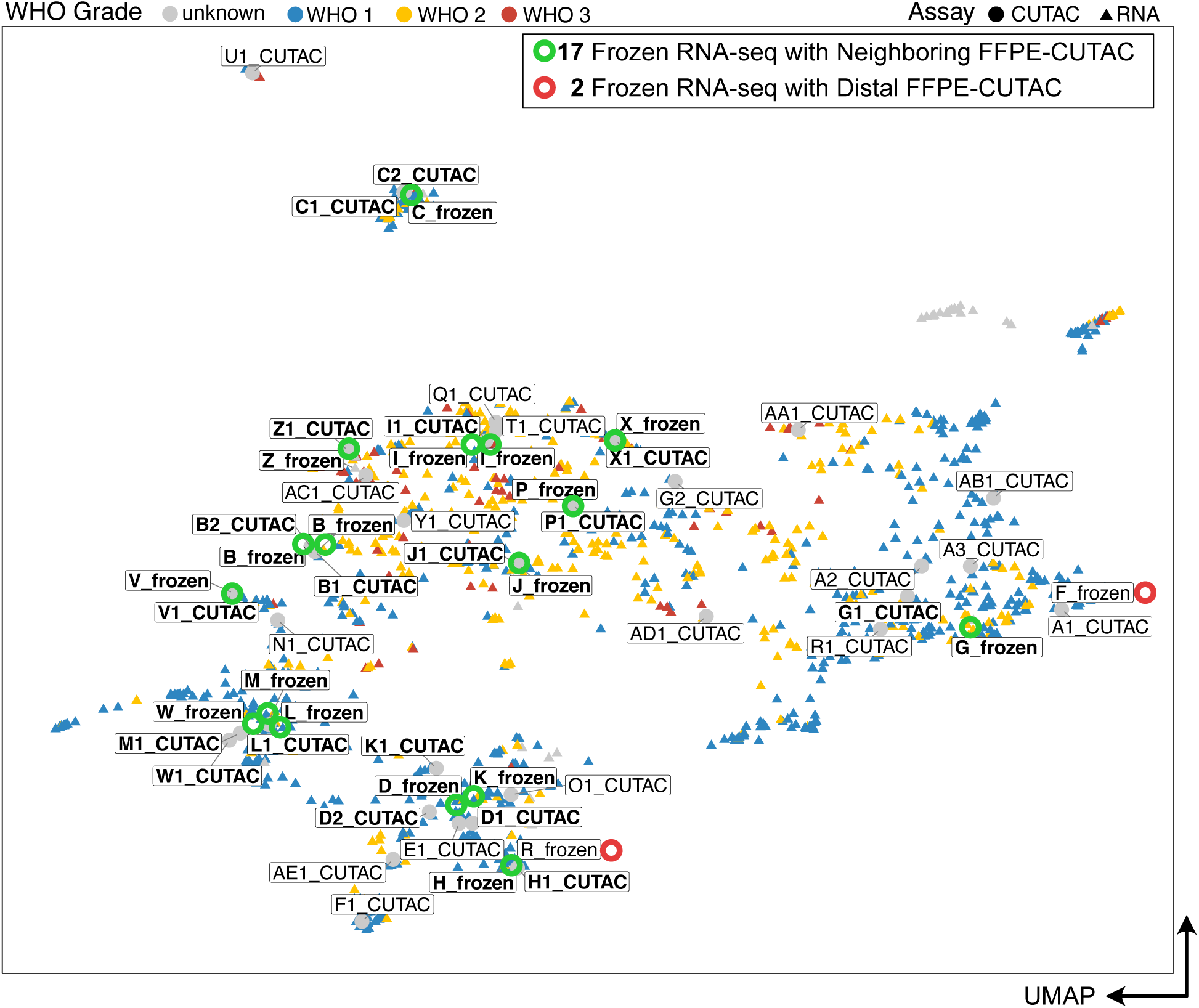
Integration of FFPE-CUTAC samples with frozen RNA-seq meningioma. Green circles mark frozen RNA-seq samples (triangles) with a close and matched FFPE-CUTAC neighbor (dots), and red circles mark RNA-seq samples without a close matching FFPE-CUTAC neighbor. Points are colored by the WHO grade.

**Fig. S13.**
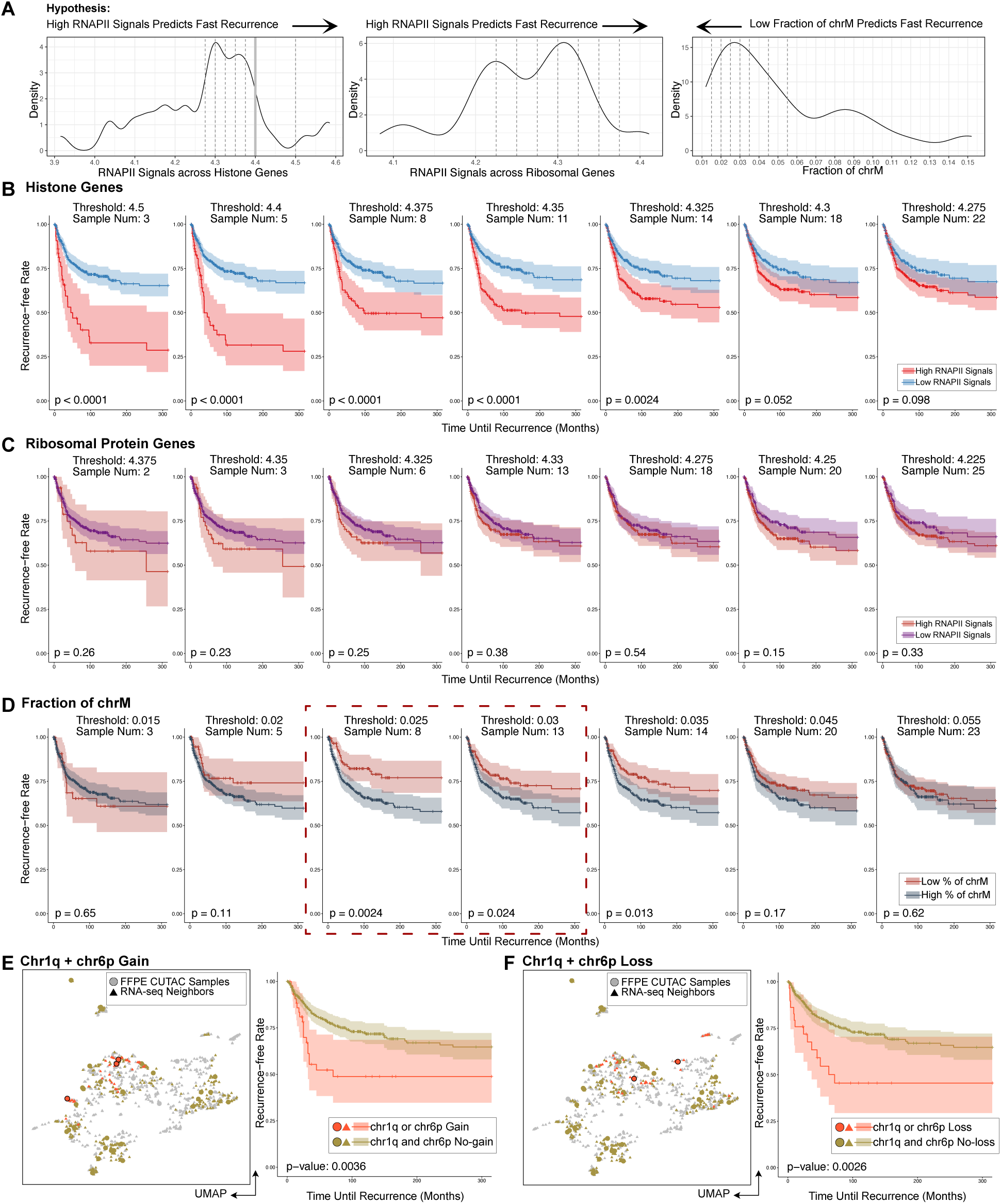
RNAPII at Histone genes predicts recurrence time of meningiomas but RNAPII at Ribosomal Protein genes or fraction of chromosome M do not. (**A**) Density distribution of the RNAPII at Histone genes (left) and Ribosomal Protein genes (middle), and the fraction of chrM (left) across 30 meningioma patients and 36 FFPE-CUTAC samples. The hypothesis is that high RNAPII signals or a low fraction of chrM predict faster recurrence. Threshold 4.4 (grey line) in the Histone gene panel is set to separate the malignant from the benign group used in Figure 4, considering most meningioma patients are benign. Dashed lines correspond to a series of separation thresholds to illustrate the survival changing pattern in B-D. (**B-D**) Kaplan-Meier (KM) curves compare the recurrence between the malignant and benign group predicted by RNAPII signals at 64 PC Histone genes (B) and Ribosomal Protein genes (C) and the fraction of chrM (D). The sample number in each panel title indicates the sample number in the predicted malignant group. The p values at the left-bottom corner of each panel are the log-rank test evaluating the separation of two survival curves. 95% confidence intervals are shown as ribbons around the curves. The dashed square highlights the significant thresholding settings for chrM, but the KM curve order is against the hypothesis that low chrM predicts malignancy. (**E-F**) Evaluation of the recurrence difference between patients with any chromosome 1q (chr1q) and chromosome 6p (chr6p) gain (E) or any loss (F), where histone genes major cluster is located on chr1q and the minor cluster is located on chr6p.

**Fig. S14.**
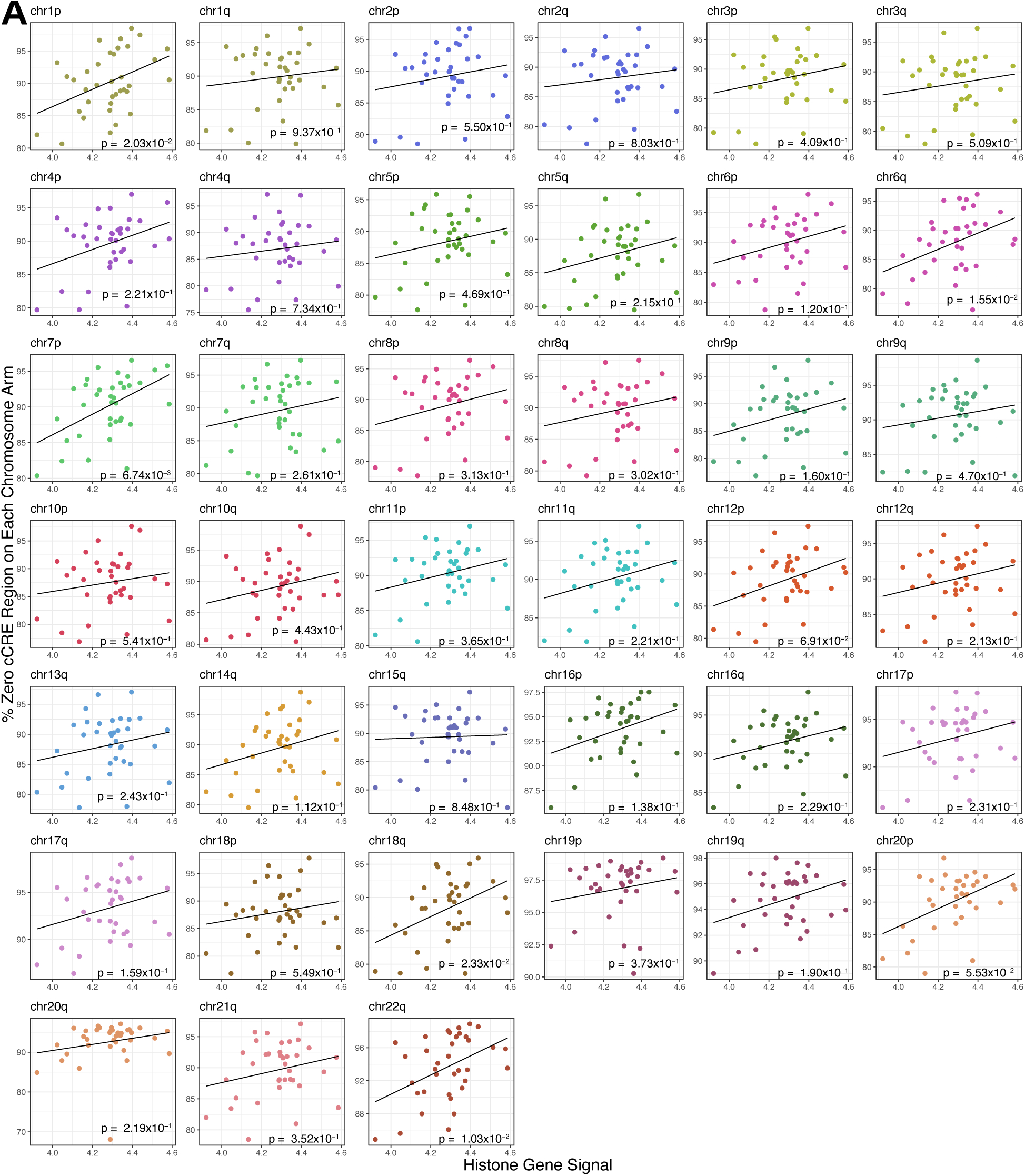
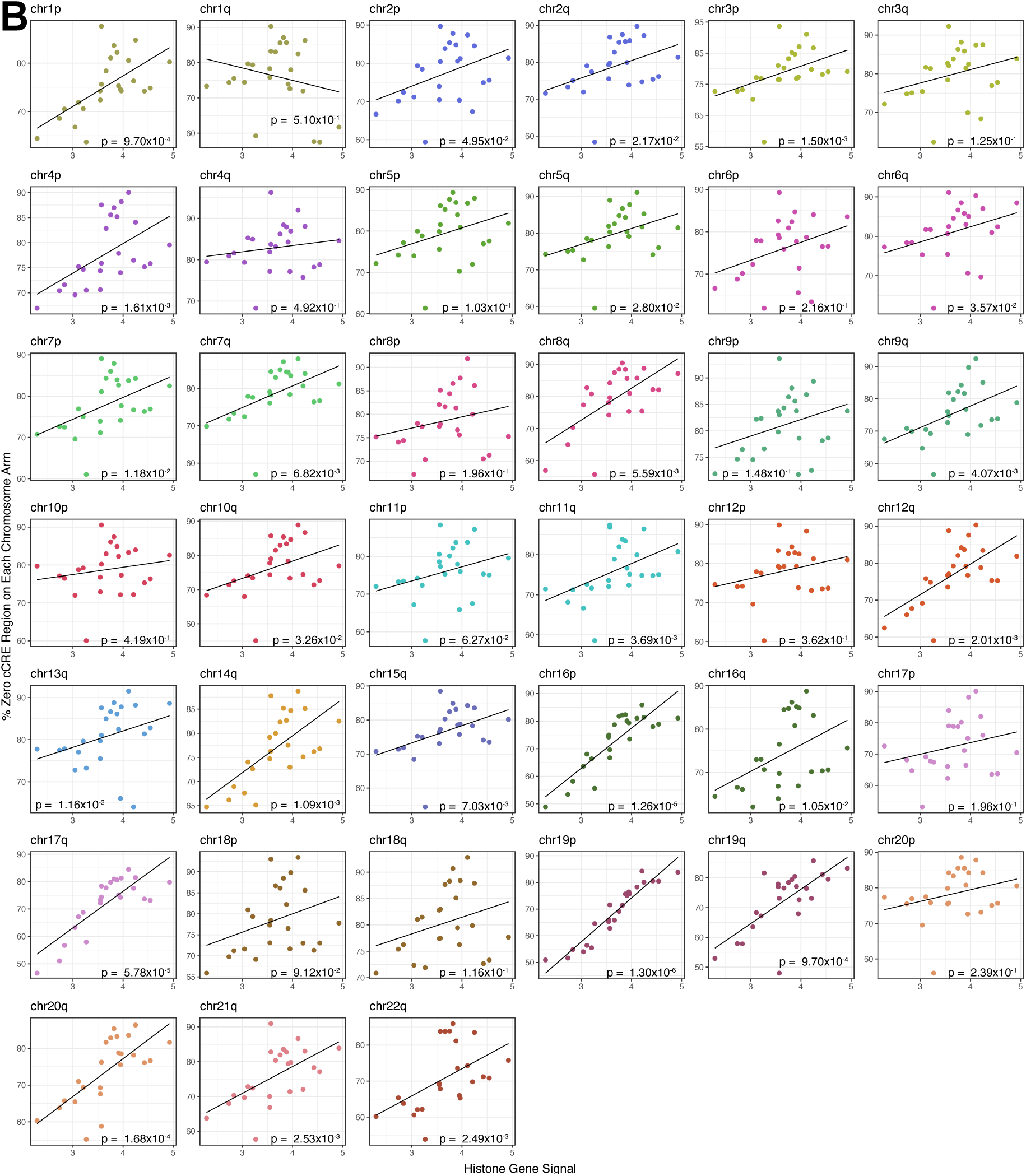
Whole-arm losses are more frequent than whole-arm gains in meningiomas as RNAPII signals increase on histone genes. To design a metric to quantify the relative frequencies of gains and losses for each of the 39 human autosomal arms (1p, 1q,…22q), we measured the percentage of cCRE regions with zero counts spanned by all cCREs on that arm for all meningiomas (**A**) and breast tumors (**B**), which yielded a single aneuploidy indicator value for each patient. Namely, if the chromosome arm loses, we should observe closer to 100% CRE regions with zero counts on that arm. We plotted this aneuploidy indicator on the *y*-axis against the aggregated RNAPII counts over the RC histone genes on the *x*-axis for that patient tumor, where the p-value for Spearman correlation is shown at the bottom of each panel. Figure 5 summarizes the correlation coefficients and significance levels on each arm.

**Fig. S15:**
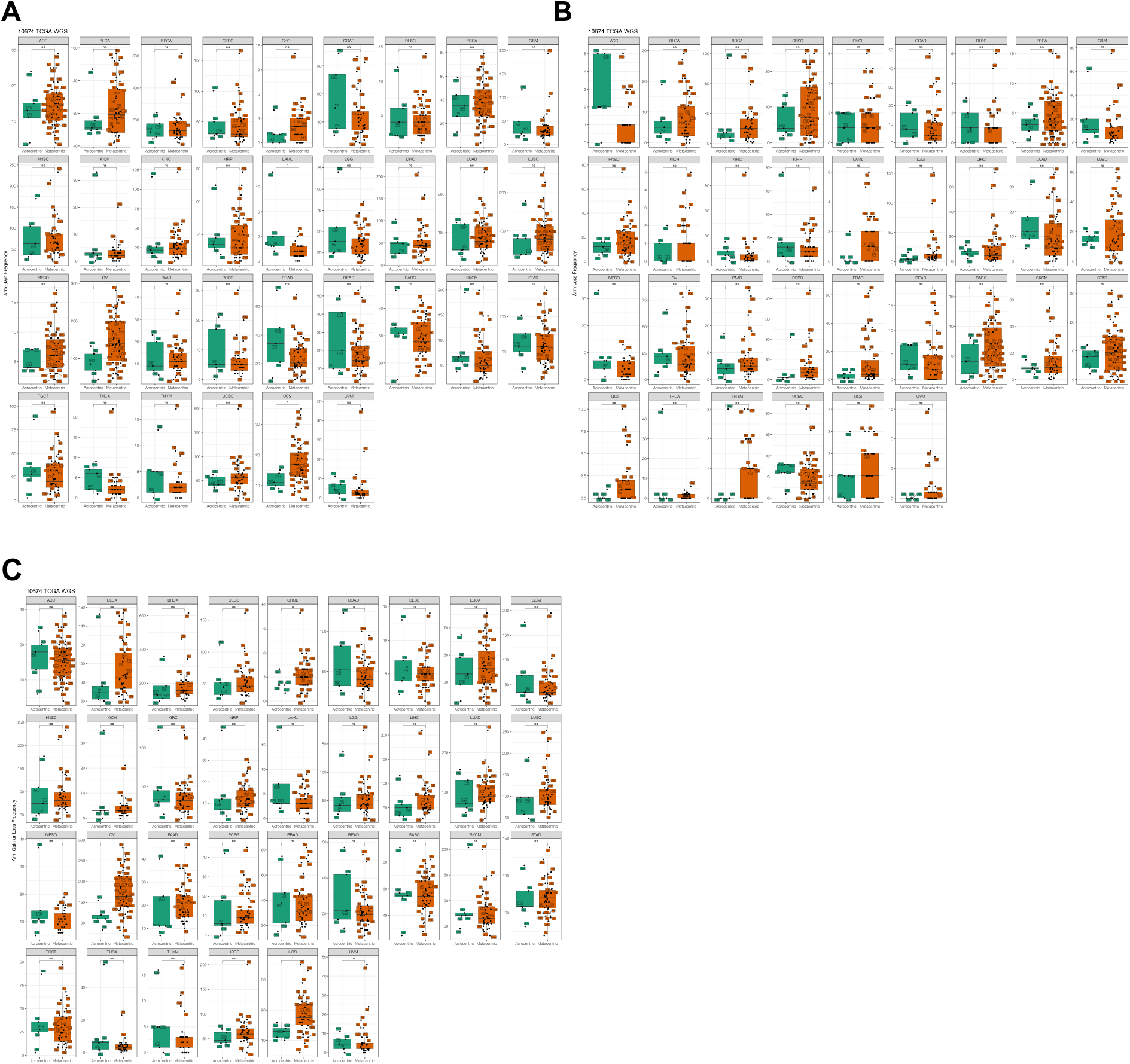
Acrocentric and metacentric whole-arm SCNAs are recovered at similar frequencies in 33 cancer types. We used whole-genome sequencing data from 10,674 cancer patients spanning 33 cancer types downloaded from The Cancer Genome Atlas (TCGA, https://portal.gdc.cancer.gov). For 32 of 33 cancer types, no significant differences are seen between acrocentrics and metacentrics in the frequencies of whole-arm SCNA gains, losses or both gains and losses (Wilcoxon rank test), except for OV, in which metacentrics are in excess at p < 0.05. Each dot represents a different autosomal chromosome arm (5 acrocentric long arms and 17 metacentrics). We inferred the chromosome arm gain or loss using allele-specific copy number analysis of tumors (<u>ASCAT)</u> (*56*) profiles for each patient. We summed the major and minor alleles of each segment and took the minimum and maximum of the copy number across segments on each chromosome arm. Any increases in the minimal allelic copy number from the diploid copy number of 2 indicate an arm gain. Similarly, any decreases in the maximum allelic copy number from the diploid copy number of 2 indicate a whole-arm loss for the corresponding autosomal arm. (**A**) Gains; (**B**) Losses; (**C**) Gains or losses.

